# *Pseudomonas aeruginosa* SPT08, a tomato endophyte, improves plant growth and controls bacterial wilt in tomato

**DOI:** 10.1101/2025.08.17.670768

**Authors:** Shubhra Jyoti Giri, Rekha Rana, Pankaj Losan Sharma, Shuhada Begum, Lukapriya Dutta, Sumi Kalita, Shuvam Bhuyan, Monika Jain, Niraj Agarawala, Prabhu B. Patil, Suvendra Kumar Ray

## Abstract

Endophytes are a class of non-pathogenic microorganisms that reside within a plant and contribute to their health. Here, we isolated several endophytes from cotyledon-stage tomato seedlings that inhibited the growth of *Ralstonia pseudosolanacearum*, the bacterial wilt pathogen of tomato. One such endophyte, *Pseudomonas aeruginosa* SPT08, protected tomato seedlings as well as grown-up plants from the wilt disease. SPT08 also improved the tomato plant height by 20% and root growth by 60% in weight. SPT08 colonization inside tomato seedlings as well as in grown-up plants was studied using green fluorescent protein to demonstrate its endophytic behavior. SPT08 exhibited twitching and swimming motility and produced extracellular enzymes such as pectinase, protease, and amylase. SPT08 tested positive for several plant growth-promoting features such as phosphate solubilization, production of siderophore, plant hormone auxin, hydrogen cyanide, and ammonia. These features were further corroborated with SPT08 whole genome sequence. SPT08 genome is 6265489 bp, with 66.59% G+C and 5786 coding genes, including type II, III, and VI protein secretion systems. The antiSMASH tool identified potential for several secondary metabolites, including antibiotics, in SPT08. This study underscores the utility of *P. aeruginosa* SPT08 bio-protection from bacterial wilt and growth promotion agent in tomato.

**Highlights:** *Pseudomonas aeruginosa* SPT08, an endophyte isolated from tomato seedlings that inhibits bacterial wilt pathogen *R. pseudosolanacearum,* can control bacterial wilt in tomato plants and enhance the height and root growth.

## 1 Introduction

*Ralstonia solanacearum* species complex, which includes different species such as *R. solanacearum*, *R. pseudosolanacearum,* and *R. syzygii,* causes a lethal wilt disease in more than 504 plant species belonging to 86 botanical families (Lowe-Power et al., 2020). The pathogen lives in the soil as a saprophyte, from where it enters the host plant through the root and then systemically colonizes the plant before wilting it (Genin, 2010; Genin and Denny, 2012; Garcia-Estrada et al., 2023). The current mode of managing the *Ralstonia pseudosolanacearum* infection includes sanitization of fields and seeds, crop rotation, application of agrochemicals, and use of various resistant varieties of seeds/crops (Wang et al., 2023; Adebayo et al., 2009; Alam and Rustgi, 2020). However, all the strategies have been found inadequate to control the wilting caused by the pathogen. Nowadays, synthetic agrochemicals are widely used to control pathogen attacks, but the infusion of chemicals to manage the disease may lead to many health-related and environmental risks. There is also evidence that the pathogen has developed resistance to various chemicals (Wang et al., 2023). Bacterial wilt is very difficult to control due to its broad host range and ability to survive in diverse environments (Nion et al., 2015; Mamphogoro et al., 2020; Benti, 2023).

Plants are inhabited by many microbes that contribute to their health. Plant-beneficial microorganisms are one of the vibrant and cost-effective ways to deal with bacterial and other plant diseases (Dinesh et al., 2015; Guo et al., 2023; Jian et al., 2024). Endophytes and plant growth-promoting rhizobacteria (PGPR) are two classes of plant-associated microbes that are widely used as biocontrol agents. Endophytes are the most promising biocontrol agents, as these microbes reside and colonize inside the plant tissues in every ecosystem without showing any disease symptoms. It is reported that endophytes show better biocontrol activity against biotic and abiotic stresses in comparison to PGPR by stimulating the host immune response, excluding the plant pathogen by niche competition or antibiosis, and some of them can enhance plant growth (Wang et al., 2021; Vinayarani and Prakash, 2018; Upreti and Thomas, 2015).

In this study, we focused on endophytic bacteria isolated from surface-sterilized tomato seedlings that exhibit the growth inhibition ability of *R. pseudosolanacearum* F1C1 in laboratory medium. One of the endophytes, *Pseudomonas aeruginosa* SPT08, also protected tomato seedlings from bacterial wilt upon mix inoculation with *R. pseudosolanacearum*. The protection was also observed in grown-up tomato plants. Interestingly, SPT08 also increases the tomato plant’s shoot height and root growth. To our knowledge, this is the first report of a native endophyte, *Pseudomonas aeruginosa* SPT08, from tomato seedlings that promotes plant height and root growth, and protects tomato plants from bacterial wilt disease caused by *Ralstonia pseudosolanacearum* F1C1. The bacterial endophyte SPT08 may act as an epigenetic factor to maintain the plant’s health and development, with a significant effect on its hosts. The dynamic tripartite interaction between the wilt pathogen-endophyte and its host plant will give an idea about controlling the wilt disease caused by *R. pseudosolanacearum*.

## 2 Materials and methods

### 2.1 Chemicals and growth media, and bacterial growth conditions

The chemicals, antibiotics, and media components of the bacterial growth medium used in this study were procured from Hi-Media, Mumbai, India, and SRL, New Mumbai, India. Glassware and plastic wares were bought from Borosil and Tarsons, Kolkata, India.

*Ralstonia pseudosolanacearum* and the endophyte strains were grown at 28 °C in BG (Bacterial growth) medium, which consists of 1 % peptone, 0.1 % casamino acid, 0.1 % yeast extract, and 0.5 % glucose (Ray et al., 2015). BG medium was supplemented with triphenyl tetrazolium chloride (0.05%). In the case of BG-Agar medium, 1.5 % agar was added to the BG medium before autoclaving. Antibiotic concentrations were used as follows: kanamycin (Kan; 50 μg/mL) and gentamycin (Gen; 50 μg/mL) when needed. *Escherichia coli* strain was grown at 37 °C in Luria-Bertani medium (Hi-Media, Mumbai).

The bacterial strains and plasmids used in this study are listed in **Table 1**.

**Table 1:**
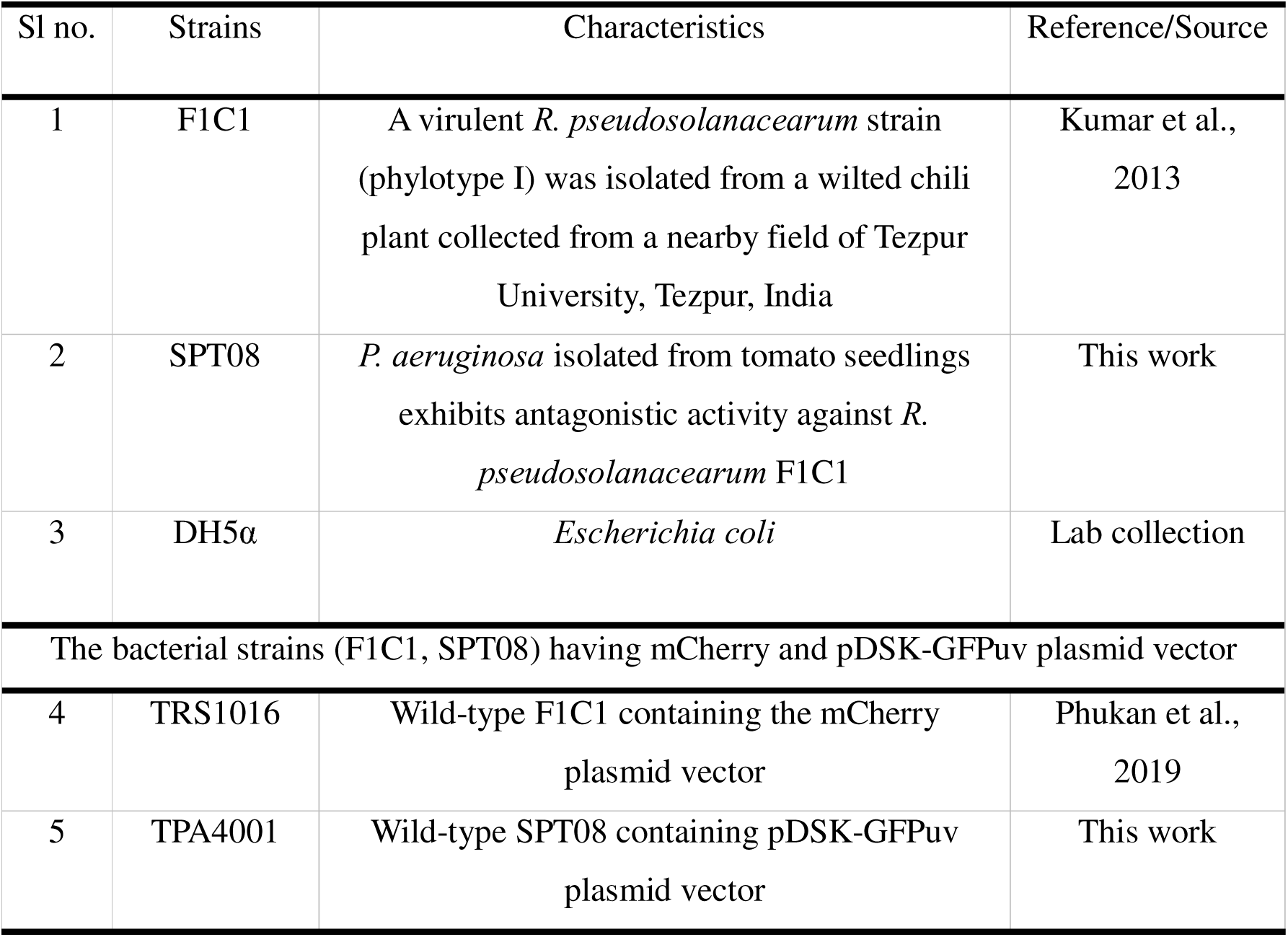
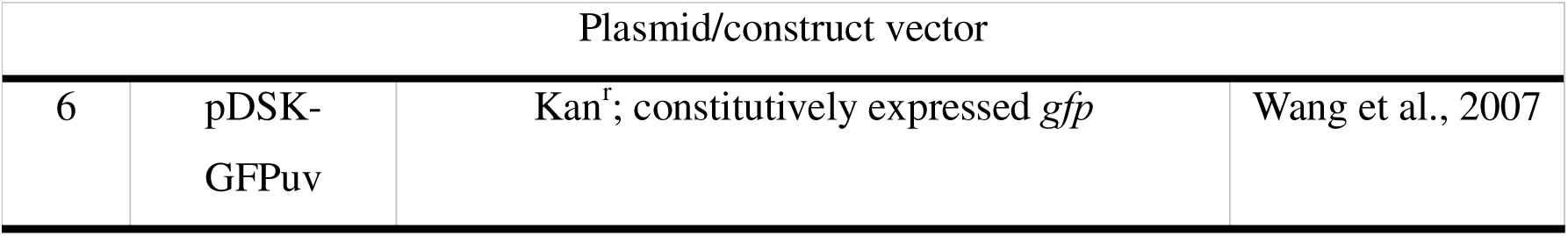
List of bacterial strains used in this study.

### 2.2 Isolation of endophytes from tomato seedlings

Tomato seeds (TO-3150, Syngenta India, Pune, Maharashtra) were collected from Tezpur local market, Tezpur -784028, Assam. The seeds were washed with sterile distilled water three times and then kept submerged for 12 h at room temperature with sterile distilled water in a glass beaker. The beaker opening face was covered by aluminium foil. The soaked seeds were then washed with sterile distilled water and then spread on wet tissue paper placed on the absorbent cotton bed. The tray was covered with a poly bag to retain the moisture inside and kept in the dark at 28 °C for two days inside a growth chamber (Orbitech, Scigenics Biotech). After the observation of white radicles sprouted from these seeds during incubation, the tray was removed from the poly bag, and the sprouted seeds were exposed directly to light and dark periods of 12 h duration each and kept at 28 °C inside the growth chamber for seven days (Kumar et al., 2017). The humidity was maintained at 75 % and the seeds were watered regularly at 12 h intervals. After seven days, ∼ 90 % seeds were found to be germinated into two-leaved cotyledon stage seedlings. These cotyledon stage seedlings were surface-sterilized using the method described in Kabyashree et al. (2020). First, the seedlings were washed with sterile distilled water repeatedly three to four times, followed by washing in Bavistin (0.04 %) solution for 2 min. Then washed with sterile distilled water three times to remove the Bavistin. Then the seedlings were washed in Mercury chloride (HgCl_2_) (0.04%) for ∼ 30 sec, followed by washing with sterile distilled water three times. The seedlings were treated with 70 % ethanol for 2 min and then washed with sterile water five times to get rid of any residual ethanol. To confirm the surface sterilization process is successful, an aliquot of water from the final rinse was plated on a BG agar plate to observe no growth of any microorganism upon incubation at 28 °C. The surface-sterilized seedlings were crushed into a homogenate using a sterile mortar and pestle. The homogenate was serially diluted (10-fold in each step) up to 10^8^-fold. A volume of 100 µL from each dilution was spread on BG-agar plates and incubated at 28 °C for 2 days. Three replica plates were spread for each dilution. Microbial colonies that appeared on the plates were re-streaked to confirm their growth and purity. The isolated strains were now taken for further characterization as well as storage at -80 °C for further studies.

### 2.3 Antagonistic activity of the endophytes against *Ralstonia pseudosolanacearum* F1C1

The antagonistic activity of the isolated putative endophytes was tested against the *R. pseudosolanacearum* using the cross-streak method as well as agar well diffusion. In the agar well diffusion method, 100 µL of *R. pseudosolanacearum* (10^9^ cfu/mL) from the saturated culture was spread on a BG-agar plate. At the center of the solid medium, using a sterile cork borer, a 4 mm diameter well was made, to which 50 µL from the saturated culture of the bacteria (10^9^ cfu/mL) was added. The plate was incubated at 28 °C for two days. Observation of a clear zone surrounding the well indicated the antagonistic activity of the endophyte. In the cross-streak method, the isolated bacteria were crossed along the diameter of the petridish, and *Ralstonia pseudosolanacearum* was streaked along the diameter in a manner that the two lines crossed at a right angle to each other. This was performed in two different patterns: (i) first, F1C1 was streaked, followed by the endophyte; (ii) first, an endophyte was streaked, followed by F1C1. The plates were incubated in static conditions at 28 °C for 48 h (Sharma, 2023).

### 2.4 Growth and generation time determination of *Pseudomonas aeruginosa* SPT08

The generation or doubling time of SPT08 was calculated as described by Bhuyan et al. (2023). Generation time of SPT08 was studied using BG and LB medium at 28 °C and 37 °C. The number of times that cells divide between the initial and final time is the number of generations for that time interval. This was calculated as 2 ^n^ = final population at (x+1) h / initial population at x h. The generation time of SPT08 was found to be ∼ 40 min at 28°C and ∼25 min at 37°C.

### 2.5 Motility study of *Pseudomonas aeruginosa* SPT08

Twitching motility was performed as described by Bhuyan et al. (2024). Twitching motility of SPT08 was studied using a glucose-free BG agar plate. The bacterial strain SPT08 was grown overnight in BG media and serially diluted to 10^4^-fold. A volume of 5 μL diluted bacterial culture was spotted on the glucose-free BG agar plate and incubated at 28 °C. The plates were observed under an inverted fluorescence microscope with a 4X objective lens (40X magnification) within a 12-18 h duration of spotting.

Swimming motility of the endophytes was studied as described by Ray et al. (2015). Swimming motility of *P. aeruginosa* was performed in a semisolid glucose-free BG agar (0.2 % agar). We have added 0.05 % tetrazolium chloride (TZC) to the medium as a growth indicator to see more clearly the bacterial growth. The soft agar was touched with a sterile toothpick dipped in the saturated culture of the bacterium, and the plates were incubated at room temperature. Swimming motility was observed by the formation of a circular zone after 12 h of incubation.

### 2.6 Characterization of *P. aeruginosa* SPT08 for extracellular enzymes

The endophytes were analyzed for the production of four extracellular enzymes, such as cellulase, pectinase, protease, and amylase.

#### 2.6.1 Cellulase activity

Cellulase enzyme production was assessed using Carboxymethyl cellulose (CMC) (Hi-Media) medium supplemented with 1.5% agar. A loop full of bacteria from freshly grown colonies was spot-inoculated at the center of the CMC agar and incubated at 28 °C for 48 h. After incubation, the plates were washed with distilled water to remove the bacteria and then flooded with 1 % Congo red solution for 48 hr at room temperature. The plates were destained with 1 M NaCl and kept overnight at room temperature. A distinct clear zone, mostly limited to the bacterial growth region, was considered positive for cellulase production (Barman et al., 2017).

#### 2.6.2 Pectinase activity

For pectinase assay, freshly grown bacterial colonies were picked up from the plate and spotted with a sterile loop on BG agar medium that was supplemented with 0.5 % pectin (Hi Media) as a sole carbon source and incubated at 28 °C for 3 days. After incubation, plates were flooded with 2 % cetyltrimethylammonium bromide (CTAB) solution for 30 min, and then the plates were washed with 0.5 M NaCl to visualize the clear zone around the bacterial growth region (Reetha et al., 2014b; Zhao et al., 2015).

#### 2.6.3 Protease activity

Protease activity of the bacterial isolate was evaluated on a skimmed milk agar (SMA) medium. The bacterial isolate was streaked on the SMA agar plate and incubated at 28 °C for 48 h. Proteolytic activity was confirmed by the formation of a zone around the bacterial colony (Devi and Thakur, 2018).

#### 2.6.4 Amylase activity

Amylase activity of the bacterial isolate was checked on starch agar plates containing beef extract 3 g/L, soluble starch 2 g/L, and agar 15 g/L. The bacterial strain was streaked in starch agar and incubated at 28 °C for 48 h. Then the plates were flooded with Lugol’s iodine solution for 20 min. The formation of a clear zone around the colony was considered positive for amylase production (Salvi et al., 2020).

### 2.7 Activity associated with Plant Growth Promotion (PGP)

The endophyte strain SPT08 was examined for five PGP traits, such as indole acetic acid (IAA), phosphate solubilization, siderophore, hydrogen cyanide (HCN), and ammonia production, using the standard protocol of Orhan, (2016); Zhao et al. (2015); Schwyn and Neilands (1987); Reetha et al. (2014); Devi and Thakur (2018), respectively.

#### 2.7.1 Indole-3 acetic acid (IAA) production

IAA production was carried out by the method described by Orhan (2016). The bacterial isolate was grown in Luria broth supplemented with 0.5 mg/mL L-tryptophan at 28 °C for 72 h in a shaking incubator. Then, 2 mL of bacterial culture was centrifuged at 4000 rpm for 10 min at 4 °C, and the cell-free supernatant was collected. For the IAA test, 1 mL of bacterial supernatant was mixed with 2 mL of Salkowski reagent (1 mL of 0.5 M FeCl_3_ in 50 mL of 35 % HClO_4_), followed by incubation in the dark for 30 min at room temperature. Production of IAA is indicated by the change in colour from pale yellow to reddish-brown or pink after dark incubation.

#### 2.7.2 Phosphate solubilization

The phosphate solubilization test was carried out by the method described by Zhao et al. (2015). Phosphate solubilization was analyzed by using phosphate solubilizing media (PSM), which was formulated by NBRIP (National Botanical Research Institute’s Phosphate Growth) medium supplemented with 15 g/L agar and 0.025 g/L bromophenol blue as an indicator. The strain was streaked on the plate and kept at 28 °C for 48 h, and the formation of a halo zone around the colony indicates the solubilization of phosphate.

#### 2.7.3 Siderophore production

Siderophore production test was carried out by the Chrome azurol S (CAS) assay as described in Schwyn and Neilands (1987). A loopful of bacterial colony was spot-inoculated on CAS agar plates and incubated at 28 °C for 48 h. The colour change of the medium from dark green to orange around the bacterial growth was considered positive for siderophore production.

#### 2.7.4 Hydrogen cyanide production

Hydrogen cyanide production was tested by using the method described in Reetha et al. (2014a). The strain was streaked on the BG agar plate supplemented with 4.4 g/L glycine. A Whatman filter paper No. 1 (110 mm) soaked in 2 % sodium carbonate in 0.5 % picric acid solution was placed on the top of the Petri plates. Plates were sealed with parafilm and incubated at 28 °C for 3 days. The change of the filter paper from yellow to brick red indicated the production of hydrogen cyanide.

#### 2.7.5 Ammonia production

Ammonia production was determined as described by Devi and Thakur (2018). The bacterial isolate was inoculated into peptone water (10 g/L) for 48 h at 28 °C in a shaking incubator. After incubation, 1 mL of Nessler’s reagent (mixture of 2.2 g HgCl_2_, 6 g KI, and 5 M NaOH) was added to the test tubes. A colour change from yellow to light brown indicates ammonia production.

### 2.8 Hypersensitive reaction (HR) assay

The hypersensitive test was done as described by Dewianty et al. (2023) and Amaria et al. (2023) with modifications. A volume of 100 µL, saturated bacterial cell suspension was infiltrated into the *Nicotiana rustica* variety of tobacco leaves using a sterile syringe on the lower surface of the leaves. The inoculated tobacco plants were observed for 24 to 48 h for the hypersensitive reaction. When HR was positive, infiltrated areas within 48 h exhibited a different decoloration and became white, paper-like texture. Sterile distilled water and *E. coli* were infiltrated as negative control, and pathogen *R. pseudosolanacearum* F1C1 was used as positive control.

### 2.9 Biocontrol activity of *P. aeruginosa* SPT08 against bacterial wilt in tomato seedlings

*R. pseudosolanacearum* F1C1 pathogenicity in tomato seedlings (*Solanum lycopersicum* var. Pusa Ruby) has been well studied by both leaf as well as root inoculation methods (Kumar et al., 2017; Singh et al., 2018). Considering the antagonistic behaviour of *P. aeruginosa* SPT08 against *R. pseudosolanacearum* F1C1 in the laboratory-grown medium, we studied the biocontrol activity of *P. aeruginosa* SPT08 against *R. pseudosolanacearum* F1C1 pathogenicity in seven-day-old tomato seedlings. The bacterial cultures were prepared as described in the method. Seedlings treated with *E. coli* (non-antagonist of RS) and without any bacterial treatment served as control groups, RS inoculated seedlings, and mixed inoculated seedlings with F1C1 and *E. coli* were taken as a positive control. The seedlings treatment sets were as follows; (a) seedlings inoculated with sterile distilled water (Control), (b) seedlings inoculated with *R. pseudosolanacearum* F1C1, (c) seedlings inoculated with endophytic strain SPT08, (d) seedlings inoculated with endophytic strain *E. coli*, (d) seedlings mixed inoculated with F1C1 and SPT08 (1:1X, 1:10X & 1:50X ratio), (e) seedlings mixed inoculated with F1C1 and *E. coli* (1:50X ratio). After inoculation, the seedling trays were kept in a growth chamber (Orbitech, India) and maintained at 28 LJC, 75 % relative humidity, and a photoperiod of 12 h. Each treatment set consisted of 40 seedlings. The experiment was repeated three times, and up to 10 days post-inoculation (DPI), each day the data for disease progression were recorded. Data was analyzed using the Kaplan-Meier and the log-rank test was used to assess the difference in the survival curves. All disease indices lower than 2 are considered as “0”, and disease indices equal to or greater than “2” are considered as “1”.

### 2.10 *Pseudomonas aeruginosa* SPT08 colonization in tomato seedlings

For the colonization study, *P. aeruginosa* SPT08 was tagged with plasmid pDSK-GFPuv, which contains one green fluorescence protein (GFP) and a kanamycin resistance gene (Wang et al., 2007). Seven-day-old tomato seedlings were inoculated with *P. aeruginosa* (SPT08/pDSK-GFPuv) by root inoculation, and seedlings inoculated with water were taken as controls. The seedlings were kept in a growth chamber at 28 °C temperatures, 75 % relative humidity, and a photoperiod of 12 h. After inoculation, from the next day onwards, some seedlings were randomly selected for colonization assessment. The seedlings’ parts, such as roots, stems, and leaves, were harvested, surface-sterilized, and examined under a fluorescence microscope (EVOS FL, Life Technologies).

### 2.11 Biocontrol activity of *P. aeruginosa* SPT08 against bacterial wilt in grown-up tomato plants

Tomato seeds (Pusa Ruby, Durga Seeds, Ayodhya, Bharat) were germinated inside the growth chamber as described previously. After four days, healthy cotyledon seedlings were carefully transferred to a sapling tray (54 cm length × 26 cm width × 4.5 cm height) filled with sterile soil and grown in greenhouse with standard growth conditions (28 °C, 16-hour photoperiod, and 75-80 % relative humidity) for ten days with regular watering to maintain optimal soil moisture. Subsequently, ten-day-old saplings were transplanted into individual pots (2 L capacity) containing 1 kg of sterile soil. The saplings were positioned centrally in the pots, ensuring the roots were fully covered and the plant was upright for uniform root development. After post-transplantation, the plants were grown for an additional 16 days under the same controlled greenhouse conditions. Throughout the experiment, plants were watered regularly and monitored daily for uniform growth. A one-month plant growth period was ensured before inoculation in all experiments, with regular watering using sterile distilled water. Tomato plants with heights ranging from 15 to 17 cm were randomly selected, and a completely randomized design was used for the experiments. The experiment was repeated three times. The treatment groups were as follows: (a) water control (C) – plants inoculated with sterile distilled water, (b) pathogen-inoculated (P) – plants inoculated with *Ralstonia pseudosolanacearum* F1C1, (c) endophyte-inoculated (E) – plants inoculated with *Pseudomonas aeruginosa* SPT08 alone, and (d) mixed-inoculated (E+P) – plants inoculated with both *R. pseudosolanacearum* F1C1 and *P. aeruginosa* SPT08. Bacterial suspensions were prepared as previously described, with final concentrations of 10LJ cfu/mL for *R. pseudosolanacearum* and 10LJ cfu/mL for *P. aeruginosa* SPT08. Each treatment involved adding 50 mL of the respective inoculum to the soil of the designated plant sets. For the co-inoculation treatment, 50 mL of a mixed suspension was applied, maintaining the aforementioned bacterial concentrations. The control plants were treated with 50 mL of sterile distilled water. Post-treatment, plants were monitored regularly for 30 days for disease progression and biocontrol efficacy by the endophyte.

*Ralstonia pseudosolanacearum* disease progression in tomato plants was studied in detail. To validate Koch’s postulates, the pathogen was re-isolated from the wilted plant and compared with the inoculated *R. pseudosolanacearum* (Abo-Elyousr et al., 2024). Further, it was confirmed by gene-specific multiplex-PCR (Kumar et al., 2013).

### 2.12 Assessment of plant growth by the endophyte *Pseudomonas aeruginosa* SPT08

To study the plant growth by the endophyte *P. aeruginosa* SPT08, the plant height was measured every 5 days for 30 days post-inoculation. After 30 days post-inoculation, plants were carefully harvested for root phenotype analysis. To minimize root damage and make soil removal easy, pots were submerged in water for 48 h. This process softened the soil, allowing for the gentle extraction of the plant root. The soil adhering to the roots was removed by proper washing, which revealed the root system that had developed. Along with root phenotype, root length, and biomass were recorded.

### 2.13 Colonization of *Pseudomonas aeruginosa* SPT08 in tomato plants

For the colonization study, *P. aeruginosa* SPT08 was tagged with plasmid pDSK-GFPuv, which contains one green fluorescence protein (GFP) and kanamycin resistance gene (Wang et al., 2007). One-month-old tomato plants were inoculated with 50 mL of *P. aeruginosa* (SPT08/pDSK-GFPuv) by adding it to the soil, and plants inoculated with water were taken as controls. The plants were kept in a growth chamber at 28 °C temperatures, 75 % relative humidity, and a photoperiod of 12 h. After ten days of inoculation, some plants were randomly selected for colonization assessment. Different sections of plant parts, such as roots, stems, and leaves, were harvested, surface-sterilized, and examined under a fluorescence microscope with 100X magnification. (EVOS FL, Life Technologies).

### 2.14 *In-planta* localization of *P. aeruginosa* SPT08 and *R. pseudosolanacearum* in tomato plants

The *in-planta* co-localization of endophyte *P. aeruginosa* SPT08 and pathogens *R. pseudosolanacearum* F1C1 was studied to know how the pathogen interacts with the endophytes inside the host plants. For the co-localization assay, mCherry-marked *R. pseudosolanacearum* (mCherryF1C1) (Monteiro et al., 2012; Capela et al., 2017; Phukan et al., 2019) and GFP-marked endophyte *P. aeruginosa* (SPT08/pDSK-GFPuv) were mixed-inoculated into the tomato plant via the soil drenching method. After 10 days of inoculation, a few plants were randomly selected to study the colonization of bacteria inside the plants. The plants were surface disinfected and observed under a fluorescence microscope at 100X magnification, and for better image overlay function was used (EVOS FL, Life Technologies).

### 2.15 Molecular Characterization

#### 2.15.1 Genome Sequencing, Genome Assembly, Annotation, and Prediction of Genes

*P. aeruginosa* SPT08 was grown overnight in PS (1% peptone, 1% Sucrose) medium at 180 rpm, 28 °C, and pelleted down at 6000 g, 5 min. Genomic DNA of SPT08 was extracted using Quick-DNA^TM^ Fungal/Bacterial Miniprep Kit (Zymo Research). Genomic DNA quality and quantity were determined using Nanodrop (DeNovix). The whole genome sequence was generated on the Illumina Nova-Seq platform at MedGenome, Bengaluru, India. Paired-end raw reads were assessed for quality check using FastQC v0.11.9 (https://www.bioinformatics.babraham.ac.uk/projects/fastqc/). Reads with a Phred score below 30 and adapter sequences were trimmed using Trim-Galore v0.6.7 (Krueger, 2015). Trimmed raw reads were *de novo* assembled using SPAdes assembler v3.13.0 (Prjibelski et al., 2020). Genome assembly parameters were estimated using Quast v5.2.0 (Gurevich et al., 2013), and genome completeness and contamination were calculated using Checkm v1.2.2 (Parks et al., 2015). Average Nucleotide Identity (ANI) value was calculated using OAU v1.2 implemented with USEARCH v11.0.667 (Lee et al., 2016). The assembled genome of SPT08 was submitted to the National Center for Biotechnology Information (NCBI) GenBank database (https://www.ncbi.nlm.nih.gov/GenBank/). The genome of SPT08 was annotated using NCBI’s Prokaryotic Genome Annotation Pipeline (PGAP) (https://www.ncbi.nlm.nih.gov/genome/annotation_prok/). For finding Plant growth-promoting genes, SPT08 genes were annotated using BlastKOALA, a KEGG (Kyoto Encyclopedia of Genes and Genomes) tool for assigning KO (KEGG Orthology) (Kanehisa et al., 2016). Biosynthetic gene clusters (BGCs) for secondary metabolites were predicted using AntiSMASH v8.0.1 (Blin et al., 2025). The presence of secretion systems was predicted using MacSyFinder v2.1.3 using the “TXSScan” model (Néron et al., 2023).

### 2.16 Statistical Analysis

All experiments were performed in triplicate, and the statistical analysis was performed using the GraphPad Prism version 8.0 software package. Analysis of variance was conducted using a one-way ANOVA test, and means were compared by Tukey’s test at a 0.05 level of confidence. The Kaplan-Meier survival curve was used for biocontrol studies in seedlings, and log-rank tests were analyzed in R software using the script provided by Bhuyan et al. (2025).

## 3 Results

### 3.1 Growth inhibition of *Ralstonia pseudosolanacearum* F1C1 by the endophyte *Pseudomonas aeruginosa* SPT08

The surface-sterilized tomato seedlings (TO-3150, Syngenta India) were crushed using a mortar and pestle, and the homogenate was dilution plated in BG glucose medium. A total of twenty-four different isolates, treated as putative endophytes, were collected based on their plate colony morphology. These endophytes were tested for their ability to inhibit the growth of *R. pseudosolanacearum* F1C1 on BG agar plates using both cross-streaking as well as agar well diffusion methods. Ten of the isolates inhibited the growth of F1C1. These ten isolates were characterized for motility, extracellular enzymes, and plant growth promotion traits (Table 2).

**Table 2:** Characterization of anti-*Ralstonia pseudosolanacearum* bacterial endophytes isolated from tomato seedlings.

| Characteristics |  | Bacterial strains |  |  |  |  |  |  |  |  |  |
| --- | --- | --- | --- | --- | --- | --- | --- | --- | --- | --- | --- |
|  |  | SPT04 | SPT05 | SPT08 | SPT11 | SPT13 | SPT14 | SPT15 | SPT17 | SPT19 | SPT20 |
| Anti-*Ralstonia* activity | Inhibit F1C1 | + | + | + | + | + | + | + | + | + | + |
| Motility assay | Twitching motility | - | + | + | + | + | + | + | + | - | - |
|  | Swimming motility | + | + | + | + | + | + | + | + | + | + |
| Extracellular enzyme production | Cellulase assay | - | - | - | - | - | - | - | - | - | - |
|  | Pectinase assay | + | + | + | + | + | + | + | + | + | + |
|  | Protease assay | + | + | + | + | + | + | + | + | + | + |
|  | Amylase assay | + | - | + | + | + | - | + | + | + | + |
| Plant growth promotion traits | Siderophore production | + | + | + | + | + | + | + | + | + | + |
|  | HCN production | - | + | + | + | + | - | + | - | - | + |
|  | Phosphate solubilization | - | + | + | + | + | + | + | + | + | + |
|  | Indole acetic acid (IAA) | + | - | + | - | - | - | - | - | - | + |
|  | Ammonia production | + | - | + | + | + | + | + | + | + | - |
| Hypersensitive response | HR response in tobacco | - | - | - | - | - | - | - | - | - | - |

Considering the maximum antagonistic activity exhibited by the isolate, SPT08, it was studied further (Fig. 1). The colony morphology of SPT08 was flat and round-shaped with a light-yellow color on the BG agar (Supplementary Fig. S1A). The bacterium has both twitching and swimming motility (Supplementary Fig. S2). It exhibited extracellular pectinase activity and no detectable level of extracellular cellulase activity. The generation time of the bacterium was found to be ∼ 40 minutes. Further, its ability to grow at 37 °C was confirmed. Its growth at 37 °C was found to be ∼ 28 mins, and the bacteria grow in BG as well as LB. The strain was initially identified as *Pseudomonas aeruginosa* from 16S rDNA sequencing. Later, whole-genome sequence-based typing confirmed that the strain SPT08 belongs to the species *Pseudomonas aeruginosa* (referred to as *P. aeruginosa* SPT08 hereafter). The scanning electron microscopy (SEM) suggested that it is a rod-shaped bacterium with a length of ∼ 1.40 µm and a width of ∼ 0.5 µm (1 µm scale bar) (Supplementary Fig. S1B).

**Fig. 1.**
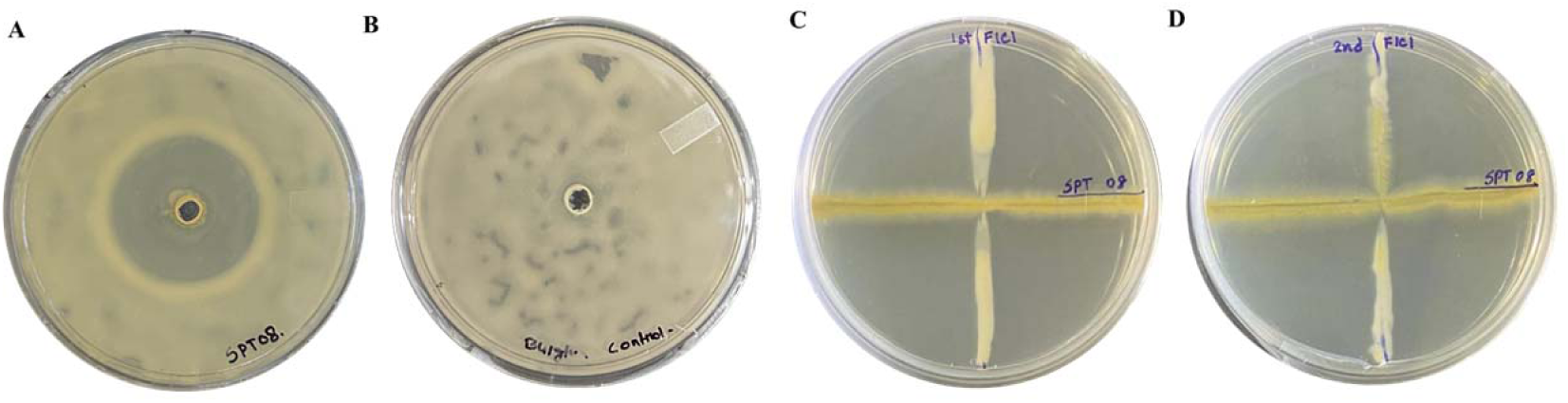
Demonstration of *Ralstonia pseudosolanacearum* F1C1 growth inhibition by *Pseudomonas aeruginosa* SPT08. The antagonistic activity of the endophyte *P. aeruginosa* SPT08 against *R. pseudosolanacearum* was assessed by agar well diffusion as well as crossed-streak methods. In the agar well diffusion, after spreading 100 μL of *R. pseudosolanacearum* culture (10^9^ cfu/mL) over the BG-agar medium in a petridish, at the center of the medium, a well was made, and then 50 μL cell suspension of *P. aeruginosa* SPT08 (10^9^ cfu/mL) was added to the well. After 48 h, a clear zone of inhibition measuring a mean diameter of 33.33 ± 0.57 mm (**plate A**) (average of three plates) was observed. This suggested that an external secretory product produced by SPT08 in the rich medium is inhibiting the growth of F1C1. In a control plate (**plate B**), instead of SPT08, 50 μL sterile distilled water was added, where no clear zone was observed. In the cross-streak method, using a sterile loop touched with a bacterial colony, first *R. pseudosolanacearum* F1C1 was streaked vertically from top (north) to bottom (south) direction, followed by the streaking of the endophyte *P. aeruginosa* SPT08 in a horizontal direction from left (west) to right (east). At the cross point, growth inhibition of F1C1 by SPT08 was observed (**Plate C**) distinctly as low growth of F1C1 was observed near the cross points in comparison to the faraway regions along the vertical line. In the reverse approach, first *P. aeruginosa* SPT08 was streaked horizontally from left to right, then *R. pseudosolanacearum* F1C1 was streaked vertically from down to up direction. The growth inhibition was observed at the cross point (**Plate D**), which was different from that of plate C. The growth of F1C1 was observed only at the down region (south direction) of the vertical line, but not at the up region (north direction) of the vertical line, instead, SPT08 was observed. This is because while streaking F1C1, it dragged cells of SPT08 from the cross point towards north, and due to the presence of both bacteria, only SPT08 growth was observed, but not that of F1C1.

### 3.2 Protection against bacterial wilt in tomato seedlings by the tomato *P. aeruginosa* SPT08

The protection ability of SPT08 against bacterial wilt in tomato seedlings was evaluated. F1C1 was mix inoculated in different proportions with SPT08 in tomato seedlings and then inoculated by leaf as well as root inoculation methods. SPT08 inoculated seedlings were similar to the water-inoculated control seedlings, either by leaf or by root, 10 days post-inoculation (DPI), which suggested that the bacterium has no negative impact on the growth of these seedlings. In the case of seedlings inoculated in the leaf with F1C1, wilting was observed on the third DPI, and the number of wilting incidences increased gradually in different DPIs. On ten DPI, the total number of wilted seedlings averaged 37 out of 40 (∼ 92.5%). Interestingly, in the case of mix inoculations with SPT08 at various concentrations, such as 1:50, 1:10, 1:1 (F1C1 to SPT08), the mean wilting incidences percentages were significantly lower, such as 17.5%, 47.0%, and 65.0%, respectively. It indicated that the higher concentration of SPT08 provided greater protection against bacterial wilt in tomato seedlings. To avoid the possibility that this protection was due to any non-specific interaction between SPT08 and F1C1, we observed that F1C1 mix inoculated with *E. coli* with a ratio of 1:50 could cause wilting only up to ∼75.0%. It is pertinent to note that *E. coli* alone inoculated in tomato seedlings resulted in no visible impact on wilting disease, like water inoculated. Similar to leaf inoculations, protection was also observed against the wilt disease in the tomato seedlings inoculated in the root by F1C1, along with SPT08, as described below. In the tomato seedlings inoculated in the root by F1C1 alone, wilting was first observed on the third DPI, and the number of wilting incidences increased gradually up to ∼ 75.0 % (on average, 30 out of 40 seedlings) by the 10 DPI. In case of F1C1 mix inoculations with SPT08 at various concentrations, such as 1:50, 1:10, and 1:1 (F1C1 to SPT08), the mean wilting incidences were only 12.5 %, 25.0 %, and 45.0%, respectively. It indicated that as the concentration of SPT08 increases, its ability to protect the tomato seedlings from bacterial wilt increases. To avoid the possibility that this protection was due to any non-specific interaction between SPT08 and F1C1, we observed that F1C1 mix inoculated with *E. coli* with a ratio of 1:50 could cause wilting up to ∼65.0%. It is pertinent to note that *E. coli* inoculated in tomato seedlings results in no wilting disease, like water inoculated. Biocontrol efficacy of SPT08 in different concentrations was observed to be significant, as shown in the Kaplan-Meier survival analysis curve (Fig. 2). The result suggested that wilting in the seedlings was effectively controlled by the endophyte *P. aeruginosa* SPT08 in tomato seedlings by the leaf as well as the root inoculations.

**Fig. 2.**
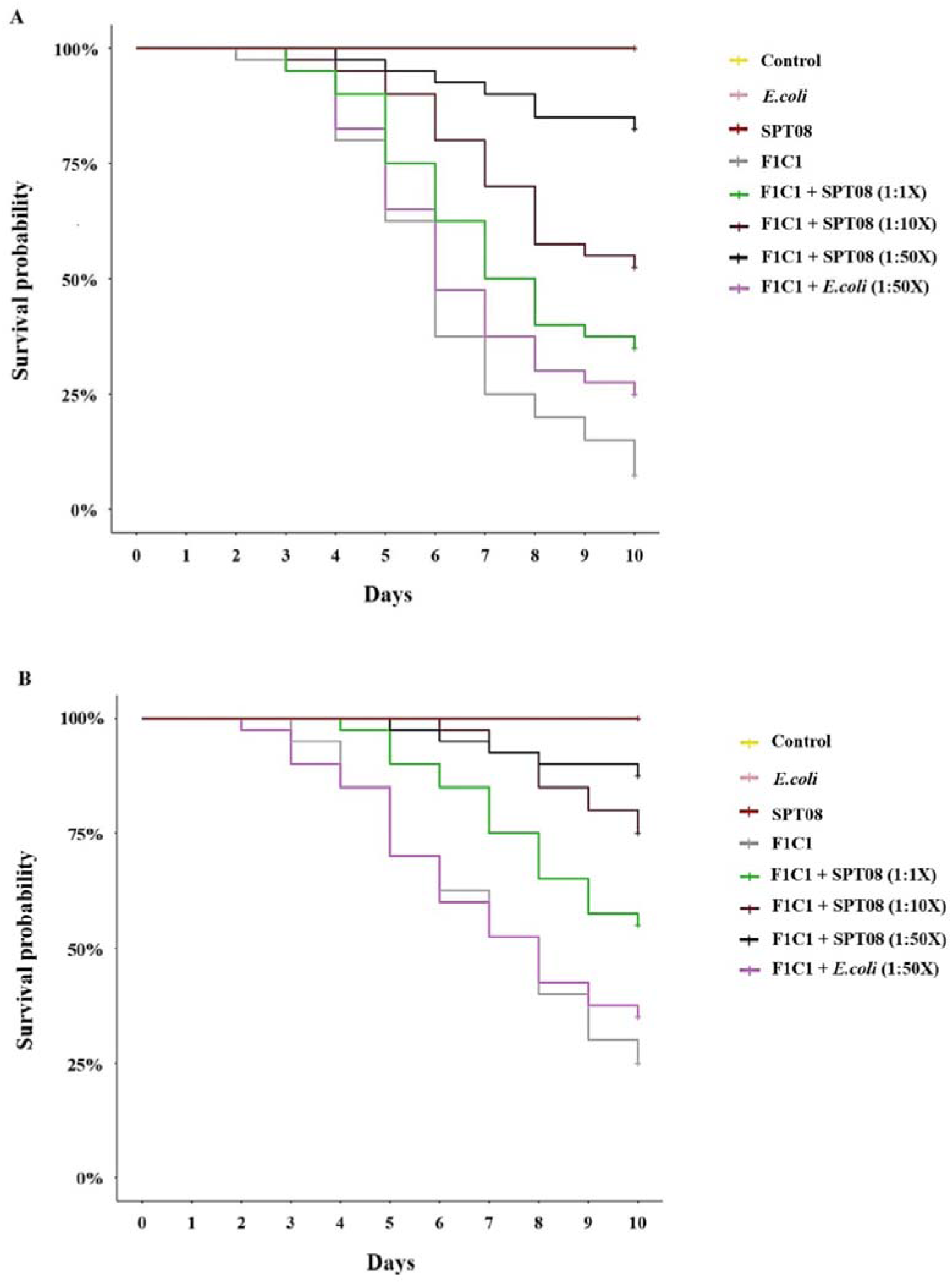
Protection against bacterial wilt in tomato seedlings by *P. aeruginosa* SPT08. Protection of bacterial wilt disease by SPT08 in tomato seedlings tested by the leaf-clip and root inoculation. Tomato seedlings were inoculated with saturated concentrations (10^9^Lcfu/mL) of bacteria. The X-axis represents days, and the Y-axis represents the percentage of survival probability. (**A**) On 10 DPI, ∼ 92.5% of seedlings were wilted upon leaf inoculation with F1C1 alone, in the case of mix inoculation of F1C1 with SPT08 (1:50, 1:10, 1:1), the mean wilting incidences percentages were significantly lower, such as 17.0% (p<0.0001), 47.0% (p<0.0001), and 65.0% (p=0.0038), respectively, F1C1 mix inoculated with *E. coli* with a ratio of 1:50 could cause wilting only up to ∼75.0% (p=0.11) in leaf inoculation. (**B**) Tomato seedlings inoculated in the root by F1C1 alone, the number of wilting incidences increased gradually up to ∼ 75.0 % by the 10 DPI, F1C1 mix inoculations with SPT08 at various concentrations, such as 1:50, 1:10, and 1:1 (F1C1 to SPT08), the mean wilting incidences were only 12.5 % (p<0.0001), 25.0 % (p<0.0001), and 45.0% (p=0.0036), respectively. F1C1 mix inoculated with *E. coli* with a ratio of 1:50 could cause wilting only up to ∼65.0% (p=0.55) in root inoculation. All data represented in the Kaplan-Meier survival analysis graph is from the three sets of experiments, and the experiments have been repeated three times.

### 3.3 *Pseudomonas aeruginosa* SPT08 colonization in tomato seedlings

SPT08 was isolated from surface-sterilized tomato seedlings with an attempt to isolate endophytes of the host plant that are antagonistic against the host pathogen. In the above, the protection ability of SPT08 against the wilt caused by F1C1 in seedlings was demonstrated. To prove that SPT08 has the ability to colonize tomato seedlings, cotyledon-stage tomato seedlings were inoculated in the root with SPT08 tagged with green fluorescence protein (GFP; SPT08/pDSKGFPuv), which is present in the plasmid pDSK-GFPuv. On 2 DPI onwards, bacterial presence was observed throughout the seedlings, such as roots, stems, and leaves, where the green fluorescence was detected in all the inoculated seedlings analyzed under the fluorescence microscope. No such fluorescence was observed in uninoculated tomato seedlings kept as controls. The uninoculated tomato seedlings served as controls, where no fluorescence signals were detected (Fig. 3). The result suggests that the *P. aeruginosa* SPT08 is successfully colonizing the tomato seedlings, proving its endophytic behaviour.

**Fig. 3.**
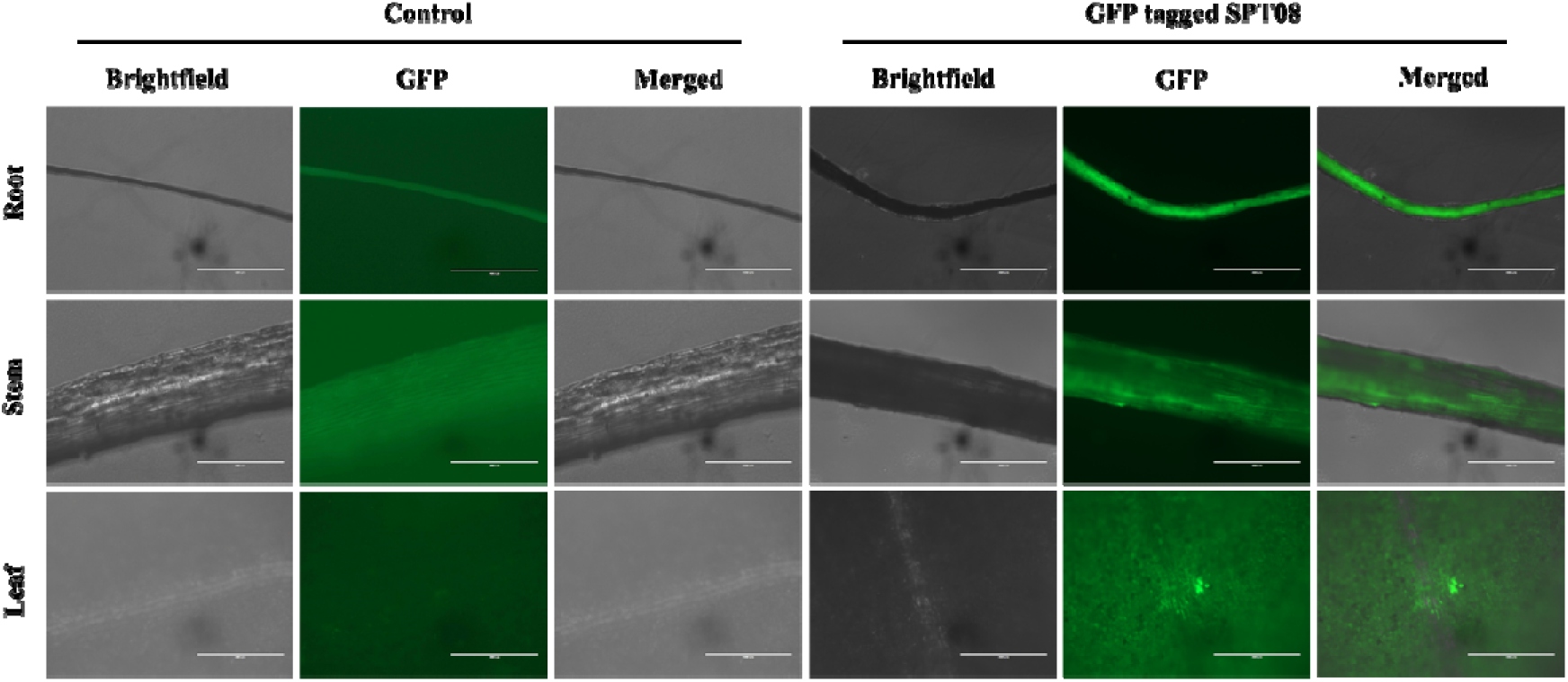
Colonization of GFP-tagged *Pseudomonas aeruginosa* SPT08 in tomato seedlings. Colonization of *P. aeruginosa* in tomato seedlings was studied at 48 h of inoculation. Tomato seedlings were inoculated through the roots with GFP-tagged *P. aeruginosa* SPT08 (10^9^ cfu/mL), and seedlings inoculated with sterile water were taken as a control. The fluorescence was observed in different parts of the seedlings, such as the root, stem, and leaf, whereas in water-inoculated control seedlings, no fluorescence was observed. Each photograph was taken using Life Technologies EVOS FL inverted fluorescence microscope with 100X magnification using the 10X objective lens with a scale bar of 400 μm.

### 3.4 *P. aeruginosa* SPT08 protection against bacterial wilt in grown-up tomato plants in the greenhouse

The biocontrol ability of *Pseudomonas aeruginosa* against the bacterial wilt in one-month-old tomato plants was studied in a greenhouse. In a set consisting of ten plants, four sets of tomato plants were inoculated separately by the soil drenching method with one set only water (C), one set with F1C1 alone (P), one set only with SPT08 alone (E), and one set with F1C1 mixed with the endophyte SPT08 (E+P). In the case of the F1C1 inoculated set, the first wilting appeared on 17 DPI. At the end of 30 DPI, all the 10 plants in the set were completely wilted (Fig. 4). It is interesting to observe that it took around thirteen days from the day the first wilting was observed till the wilting of the last plant in the set. The wilting pattern in seedlings and in grown-up plants might not be identical. No wilting was observed in the other three sets till 30 DPI. One of the wilted plants was tested for the ooze streaming test, and the presence of F1C1 in the ooze was confirmed by phylotype-specific multiplex PCR, twitching motility. Even till 40 DPI, no wilting could be noticed in the case of plants inoculated with F1C1 and SPT08. The result indicated that SPT08 has potential for biocontrol against bacterial wilt. The biocontrol ability of SPT08 was successfully observed in two other different experiments performed further in the greenhouse (Supplementary Fig. S3).

**Fig. 4.**
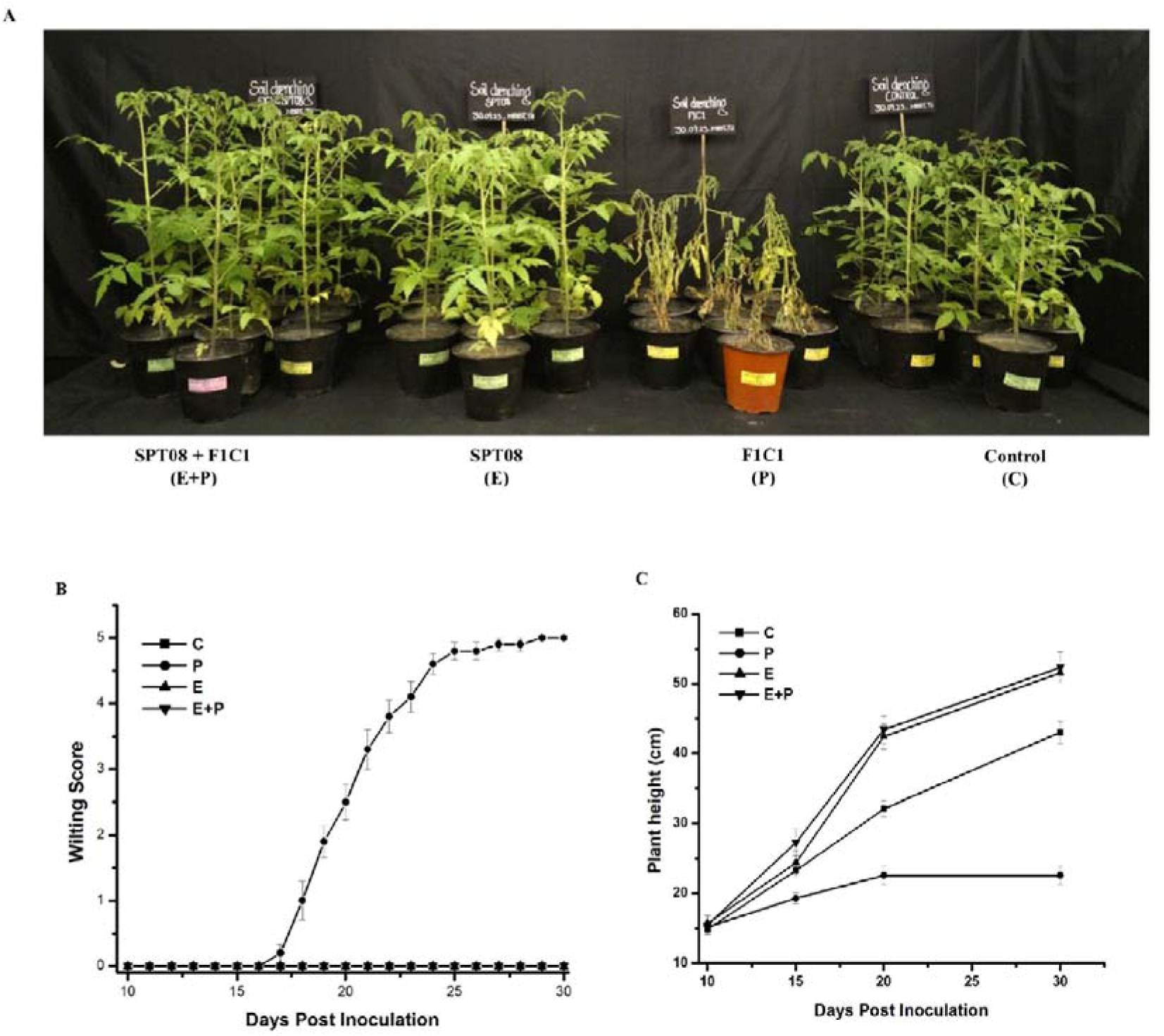
*P. aeruginosa* SPT08 protection against bacterial wilt in grown-up tomato plants in the greenhouse. Bacterial wilt protection in grown-up tomato plants by *Pseudomonas aeruginosa* SPT08 under greenhouse conditions (**A**). Four groups of one-month-old tomato plants, each with 10 plants, were inoculated by soil drenching. The inoculated plant sets are as follows: water control (C), *R. pseudosolanacearum* F1C1-pathogen inoculated (P), *P. aeruginosa* SPT08-endophyte inoculated (E), and *P. aeruginosa* SPT08 and *R. pseudosolanacearum* F1C1-mixed inoculated (E+P). The photograph was taken on 30 dpi, which represents the plant protection ability by the endophyte *P. aeruginosa* SPT08 in mix-inoculated (E+P) plants, and further, the plants inoculated with SPT08 (E) showed no symptoms, and the plants were healthy till the end of the experiment. (**B**) The graph shows the combined data of wilting disease progression studied in inoculated tomato plants after the first symptoms appeared till 30 days post-inoculation. The X-axis represents days post inoculation (DPI), and the Y-axis represents the wilting scores (0-5). Error bars shown are the standard deviation values. The wilting score was recorded using 0-5 disease index scale, where 0=no visible wilting symptoms, 1= partial wilting (one leaf or branch), 2= two leaves or branches wilted, 3= 4 to 5 leaves wilted, 4= all leaves or branches started to wilt, and 5= completely wilted/ dead/ collapsed. The wilting patterns of F1C1 inoculated plants were compared with SPT08 (E), mix-inoculated SPT08 and F1C1 (E+P), and water-inoculated control (C) plants under greenhouse conditions. (**C**) The graph shows the plant height of four sets of inoculated tomato plants at different days post-inoculation. Each treatment group consisted of 10 plants, and the experiment was repeated three times. Error bars represent standard errors.

The colonization ability of the endophyte was studied in grown-up plants by inoculating with SPT08 tagged with green fluorescent protein (GFP; SPT08/pDSKGFPuv). After 10 DPI, the presence of GFP was observed in root, stem, and leaf parts of the plant in all five inoculated plants analysed. There was no detectable level of GFP in the uninoculated tomato plants that served as controls. The result indicated that colonization of the endophyte *P. aeruginosa* SPT08 in tomato plants is systemic. We also studied the localization of SPT08 tagged with GFP and F1C1 tagged with mCherry after mix inoculation in tomato plants by the soil drenching method. Several tomato plants were analyzed after 10 DPI, where both GFP and mCherry were detected in all the plants studied (Fig. 5). It indicated that F1C1 could colonize tomato plants in the presence of SPT08 when both are co-inoculated. However, the absence of wilting in the co-inoculated plants either might be due to the low concentration of F1C1 inside the tomato plant in the presence of SPT08 and/or due to any defense response of tomato elicited because of SPT08 inside it that suppresses disease causing ability of F1C1.

**Fig. 5.**
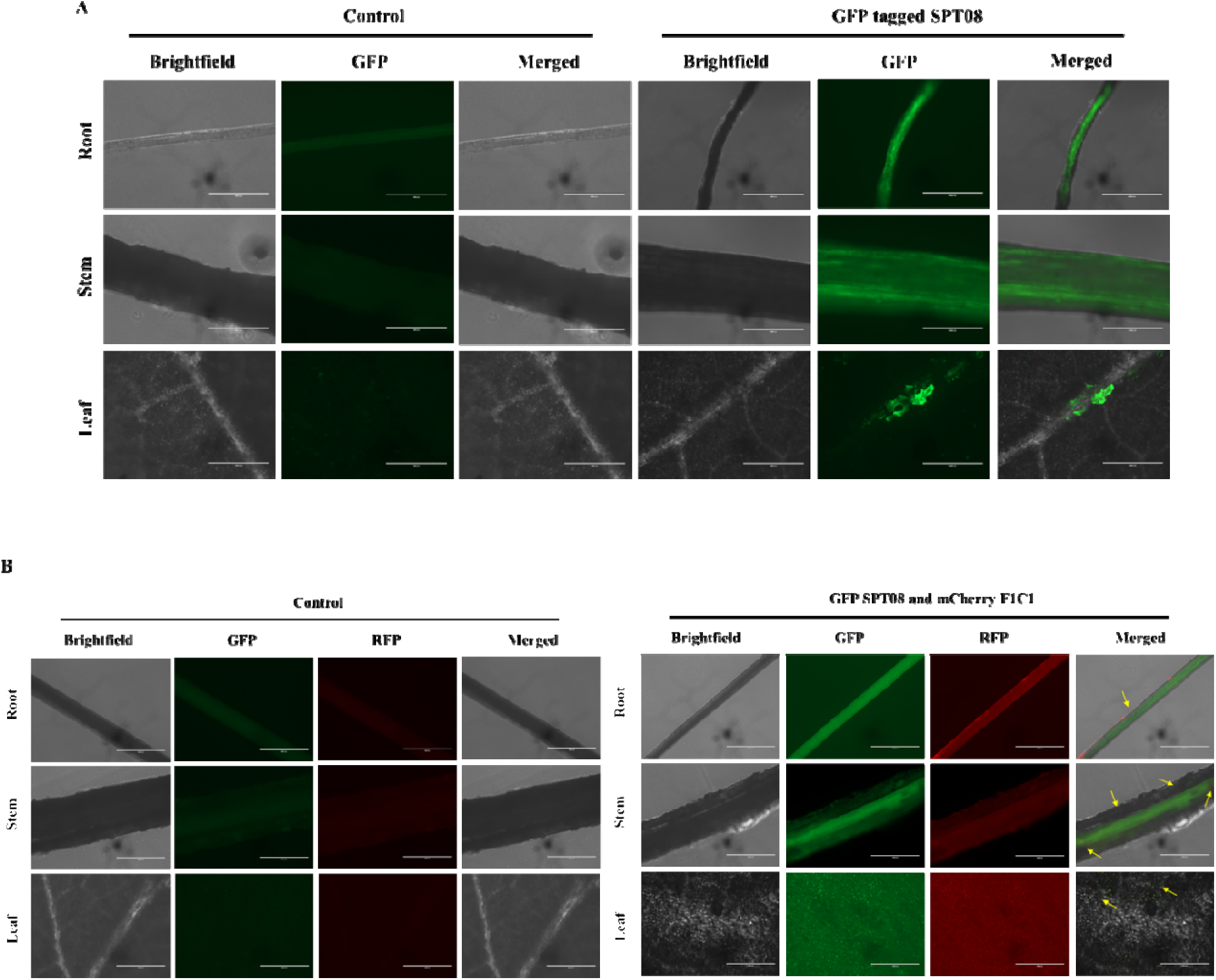
Colonization and co-localization in grown-up tomato plants. Colonization of *P. aeruginosa* SPT08 in the grown-up tomato plants was studied by inoculating GFP-tagged SPT08. After 10 days of inoculation, the colonization was observed in plants roots, stems, and leaves by fluorescence microscope (Life Technologies EVOS FL) with 100X magnification (scale bar 400 μm). (**A**) Fluorescence micrographs of root, stem, and leaves from tomato plants inoculated with water control (left), where no fluorescence was observed, whereas plants inoculated with GFP-tagged *P. aeruginosa* SPT08 showed fluorescence in the root, stem, and leaf. (**B**) Localization of GFP-tagged *P. aeruginosa* SPT08 (TPA4001) and mCherry-tagged *R. pseudosolanacearum* (TRS1016), fluorescence of both GFP SPT08 and mCherry F1C1 was observed in the root, stem, and leaf region (right), and no fluorescence was observed in the water-inoculated control plants (left).

### 3.5 *Pseudomonas aeruginosa* SPT08 increases the height of tomato plant and its root growth

Interestingly, tomato plants inoculated with either SPT08 alone or in combination with SPT08 and F1C1 were noticed to be taller than the control set of plants inoculated with water. After 30 DPI, the average plant height recorded for the control set inoculated with water was: 43.0 ± 5.01 cm, whereas for the set inoculated with SPT08 it was 51.5 ± 3.20 cm, and the set mix-inoculated with SPT08 with F1C1 was 52.3 ± 6.97 cm (Fig. 6). The plants treated with SPT08 were 19.76% taller (p=0.0031), and the plants inoculated with SPT08 with F1C1 were 21.62% taller (p=0.0011) than the control set. It is pertinent to note that the height of the plants inoculated with F1C1 was only 29.51 ± 5.9 cm, as the plants wilted by 30 DPI. The increase in tomato plant height inoculated with either SPT08 or inoculated in combination with SPT08 and F1C1 in comparison to the control tomato plants inoculated with only water was also observed in the two other times the experiment was repeated in the greenhouse (Table S1, Fig. S4).

**Fig. 6.**
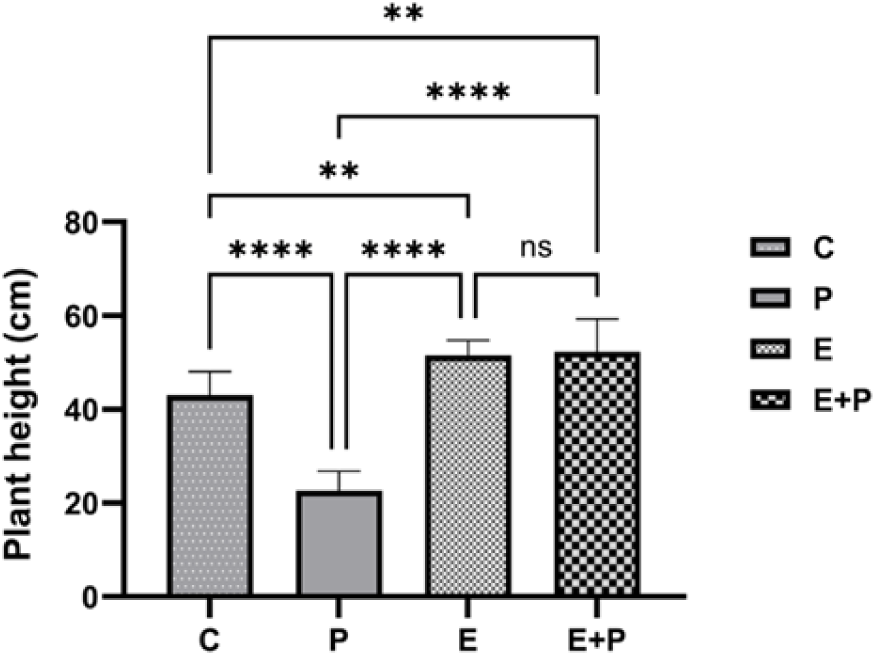
*Pseudomonas aeruginosa* SPT08 promotes the growth of the tomato plant height. Endophyte-inoculated plants were observed to increase in plant height as compared to the water-inoculated control plants after 30 days post-inoculation, as shown in the graph. Plant inoculated with sterile water represented as C, P represents pathogen F1C1 inoculated plants, E represents endophyte SPT08 inoculated plants, and E+P represents plants inoculated with SPT08 and F1C1. Plant height presented here is the average of 10 plants per treatment, and the experiment has been repeated three times. Vertical bars represent the mean value, and error bars represent the standard deviation. Bars indicate significant differences as defined by one-way ANOVA with Tukey’s HSD test (pL<L0.05). * significance at 0.05 (*p* < 0.05); *** significance at 0.001 (*p* < 0.001) and **** significance at 0.0001 (*p* < 0.0001), and ns represent non-significance.

Any morphological changes occurring in roots during plant growth are not possible to observe, as roots are covered by soil and are invisible to the outside. But, during our regular visit to the greenhouse for observing the plants inoculated with SPT08, F1C1, etc., surprisingly, one day we noticed a prominent outgrowth of roots underneath the base of the pots in the case of tomato plants inoculated with SPT08 and SPT08 with F1C1 (Fig. 7). This growth was not observed so well in the case of control tomato plants. It indicated a possibility of higher growth of roots occurring in these two sets of plants in comparison to the control set of plants. So, after the completion of the experiments, on 40 DPI, we carefully extracted the roots of these plants from the soil to which they were tightly tethered by keeping them submerged in water for 48 hours, followed by repeatedly washing under tap water. The root growth was then measured by their wet weight and dry weight (Table 3). The average fresh root weight for the control set inoculated with water was 7.13 ± 1.66 g, whereas for the set inoculated with SPT08 was 12.90 ± 2.84 g, and the set inoculated with SPT08 and F1C1 was 16.06 ± 3.48 g (Fig. 8A). The fresh root mass in plants set inoculated with SPT08 was 80.92 % higher (p<0.0001), and the plants inoculated with SPT08 and F1C1 were 109.57 % higher (p<0.0001) as compared to plants inoculated with water in the control set. The dry root mass in case of the control set inoculated with water was 0.94 ± 0.2 g, whereas in the case of SPT08 inoculated plant set was 1.48 ± 0.26 g, and the plant set inoculated by SPT08 with F1C1 was 1.97 ± 0.54 g (Fig. 8B). The dry root weight in case of plants treated with SPT08 were 57.44 % (p=0.0114) more, in the case of the plants inoculated with SPT08 and F1C1 was 109.57 % (p<0.0001) more than the control plant set. It is pertinent to note that the roots collected from the wilted plants, which were inoculated with F1C1, weighed only 1.14 ± 0.91 g as fresh weight and 0.62 ± 0.26 g as dry weight. This experiment indicated that SPT08 increases tomato plant root growth. The increase in tomato plant root mass inoculated with SPT08 or SPT08 and F1C1 in comparison to the control tomato plants inoculated with water was also observed in the two other times the experiment was repeated in the greenhouse (Supplementary Table S1, Fig. S6).

**Fig. 7.**
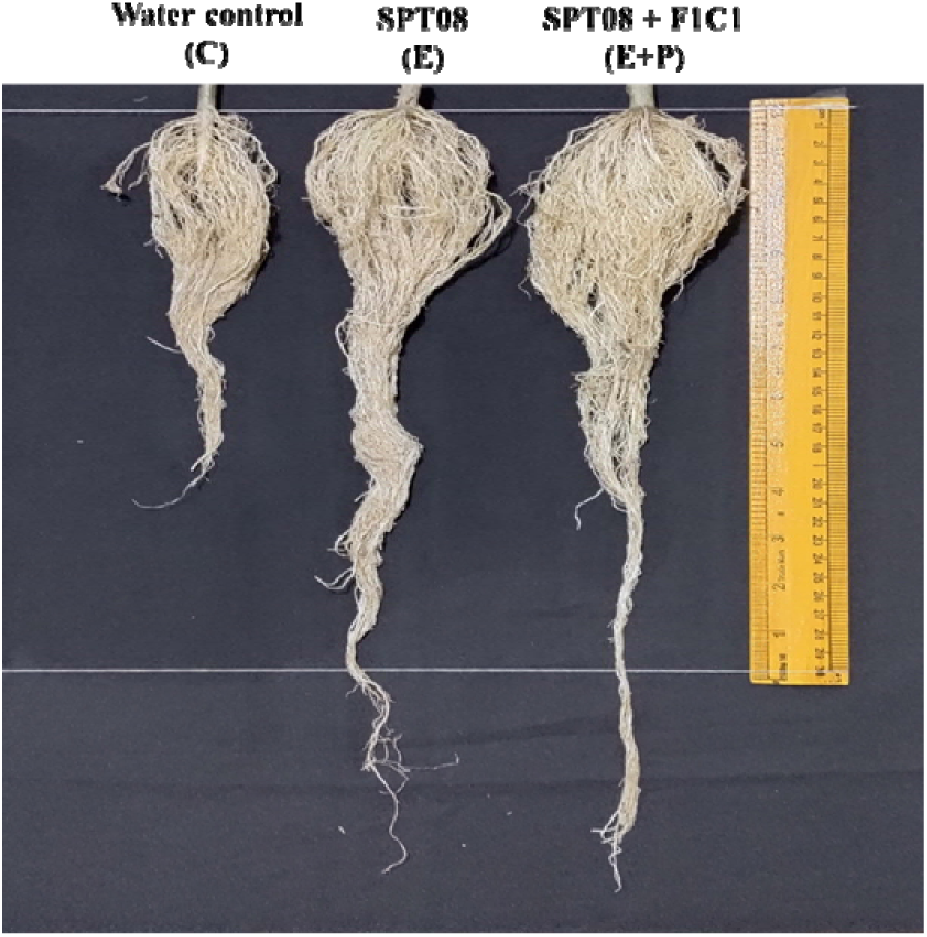
*Pseudomonas aeruginosa* SPT08 changes the root morphology of the tomato plant. Change in root architecture (RA) of tomato plants inoculated with *P. aeruginosa* SPT08. Plants inoculated with sterile water (C), endophyte *P. aeruginosa* SPT08 alone (E), and mix-inoculated with SPT08 and F1C1 (E+P), were analyzed for root growth at 40 days post-inoculation. Tomato plant roots inoculated by endophyte SPT08 (in sets E and E+P) were observed to be highly branched and thick, with increased root hair as compared to water control (C) plants. The experiment has been repeated three times, and we have observed the change in root phenotype in SPT08 inoculated plants.

**Fig. 8.**
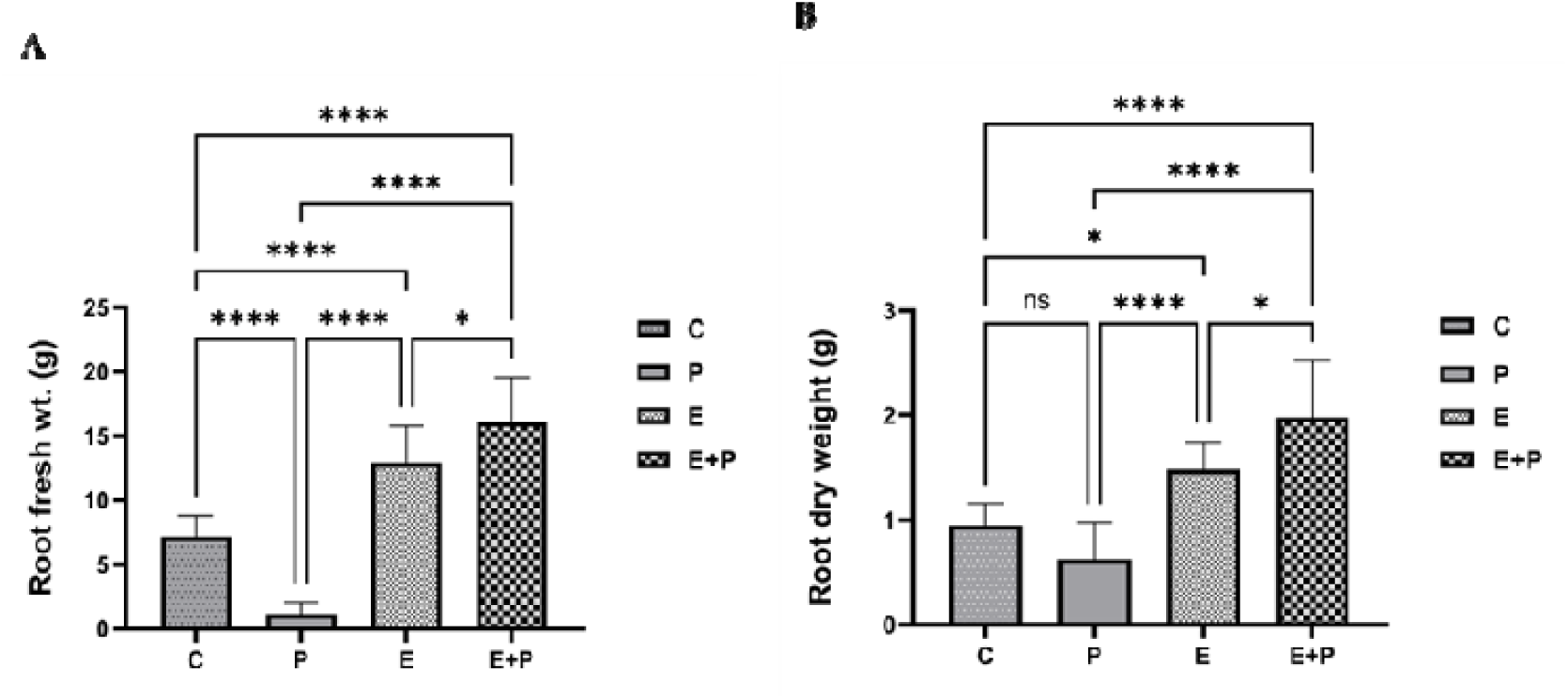
Increase in root fresh weight and dry weight by endophyte *P. aeruginosa* SPT08. Effect of endophyte SPT08 on tomato plant root growth parameters at 40 days post-inoculation. The root fresh weight (**A**) and dry weight (**B**) of SPT08 inoculated plants were compared with the water-inoculated control plants. The root fresh weight of all the tomato plants was taken on 40 dpi, whereas the root dry weight was recorded on 41 dpi. The plant root weights presented here are the average of 10 plants per treatment, and the experiment has been repeated three times. Vertical bars represent the mean value, and error bars represent the standard deviation. Bars indicate significant differences as defined by one-way ANOVA with Tukey’s HSD test (pL<L0.05). * significance at 0.05 (*p* < 0.05); *** significance at 0.001 (*p* < 0.001) and **** significance at 0.0001 (*p* < 0.0001), and ns represent non-significance.

**Table 3:** Root mass and length of tomato plants at 40 DPI (Days post inoculation).

| **Root Growth Parameters** | **Control (C)** | **F1C1 (P)** | **SPT08 (E)** | **SPT08+F1C1 (E+P)** |
| --- | --- | --- | --- | --- |
| Root fresh wt. (g) | 7.13±1.66 | 1.14±0.91 | 12.9±2.84 | 16.06±3.48 |
| Root dry wt. (g) | 0.94±0.2 | 0.62±0.34 | 1.48±0.26 | 1.97±0.54 |
| Root length (cm) | 24.3±5.31 | 15.1±4.43 | 35.95±9.49 | 32.4±4.83 |

We have also measured the root length and average root length measured for the control set inoculated with water was 24.3 ± 5.31 cm, whereas for the set inoculated with SPT08 was 39.95 ± 9.49 cm, and the set inoculated with SPT08 with F1C1 was 32.40 ± 4.83 cm (Fig. 9). The root length of the tomato plants inoculated with SPT08 and SPT08 with F1C1 was observed to exhibit significantly higher growth, increasing by 64.40 % (p=0.0012) and 33.33 % (p=0.0345), respectively. Root length of tomato plants inoculated with F1C1 was 18.103 ± 3.9 cm. The increase in tomato root length inoculated with SPT08 or SPT08 and F1C1 in comparison to the control tomato plants inoculated with water was also observed in the two other times the experiment was repeated in the greenhouse (Supplementary Table S1, Fig. S7).

**Fig. 9.**
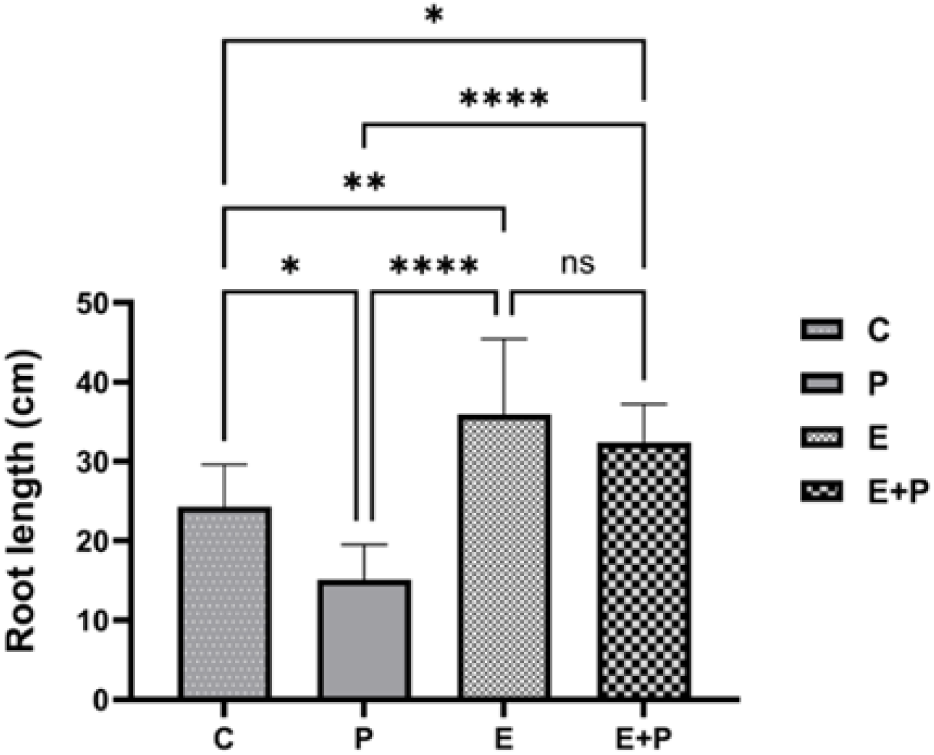
Longitudinal root length of tomato plants inoculated with *P. aeruginosa* SPT08. Root length of tomato plants inoculated with endophyte SPT08 (E) and mix inoculated with SPT08 and F1C1 (E+P), was compared with control (C) plants inoculated with sterile water after 40 dpi. The root length of endophyte-inoculated plants was observed to be higher than that of water-inoculated tomato plants. Error bars represent the mean ± SD (n=10) and bars indicate significant differences as detected by one-way ANOVA with Tukey’s HSD test (pL<L0.05). * Significance at 0.05 (*p* < 0.05); *** significance at 0.001 (*p* < 0.001) and **** significance at 0.0001 (*p* < 0.0001), ns represent non-significance.

Not only the growth morphology, but with careful observation, the roots of the plants inoculated with SPT08, as well as SPT08 with F1C1, were observed to be highly branched and thick, with increased root hairs. It is pertinent to note that several rhizospheric bacteria are known to affect plant root architecture and morphology. But this is the first report of an endophyte is protecting against bacterial wilt and is involved in changing the root architecture. The change in root architecture of tomato plants by SPT08 was also observed in two other independent experiments under greenhouse conditions (Supplementary Fig. S5).

### 3.6 Extracellular enzyme production and plant-growth promotion traits by *P. aeruginosa* SPT08

*P. aeruginosa* SPT08 was qualitatively screened for extracellular enzyme production using the agar plate method. The endophyte was found to be positive for pectinase, protease, and amylase activity. Additionally, SPT08 does not exhibit cellulase activity, while F1C1 was included as a positive control (data not shown).

*P. aeruginosa* SPT08 was qualitatively analyzed for its plant growth promotion (PGP) traits. SPT08 showed positive response for phosphate solubilization, production of indole acetic acid (IAA), siderophore, hydrogen cyanide (HCN), and ammonia (Fig. 10 and Table 4). Indole acetic acid production by the endophytic strain SPT08 was confirmed by a light pink color change, indicating positive for IAA synthesis. Siderophore production was observed by the formation of an orange halo zone on Chrome Azurol S agar after 24 h of incubation. Hydrogen cyanide production was confirmed by a color change of filter paper from yellow to brown, with an uninoculated plate serving as a negative control showing no HCN production. Phosphate solubilization was confirmed by the formation clear zone around the colony, confirming the potential phosphate solubilization activity, and ammonia production was confirmed by a change in yellow to brown color. Ammonium sulfate was taken as a positive control, and uninoculated media as a negative control. These findings highlight SPT08’s potential to enhance plant growth through diverse mechanisms.

**Fig. 10.**
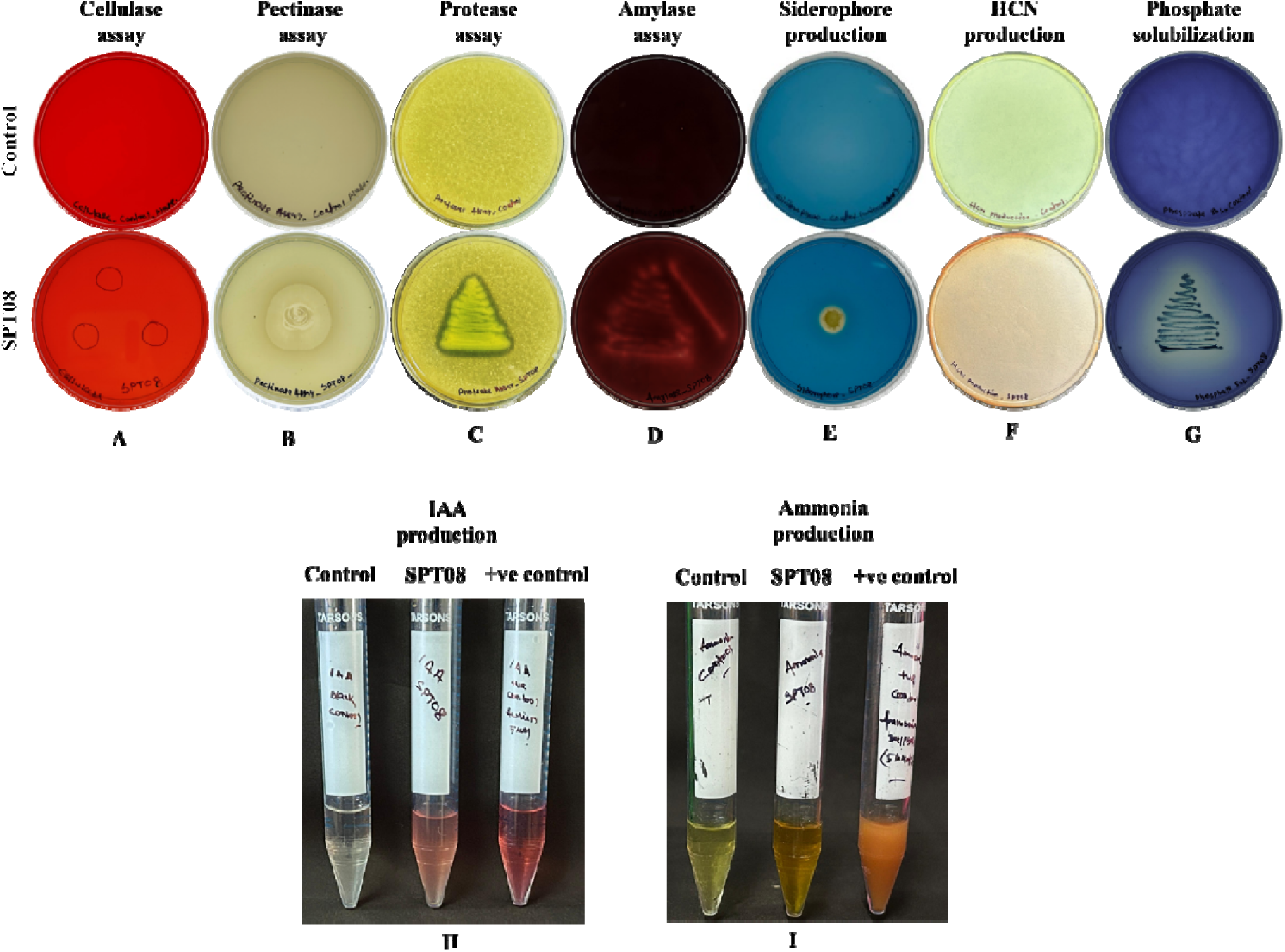
Extracellular enzyme production and plant-growth promotion traits by *P. aeruginosa* SPT08. The images represent qualitative enzyme production and PGP traits by *P. aeruginosa* SPT08. Extracellular enzymes assessed for SPT08, such as cellulase (**A**), pectinase (**B**), protease (**C**), and amylase (**D**). SPT08 was found to be positive for all enzymes assessed except cellulase. The plant growth-promotion traits by SPT08 were found to be positive for siderophore production (**E**), hydrogen cyanide production (**F**), phosphate solubilization (**G**), indole acetic acid (IAA) production (**H**), and ammonia production (**I**), in laboratory-grown respective mediums.

**Table 4:** Characterization of *P. aeruginosa* SPT08 for enzymes and plant growth-promotion traits.

| **Characteristics** | **Activity** | **Present/absent** | **Remarks** |
| --- | --- | --- | --- |
| Motility assay | Twitching motility | + | Genes encoding Type IV pili present |
|  | Swimming motility | + | Flagella |
| Extracellular enzyme production | Cellulase assay | - | No genes for cellulase |
|  | Pectinase assay | + | - |
|  | Protease assay | + | Genes are present |
|  | Amylase assay | + | Genes are present |
| Plant growth promotion traits | Siderophore production | + | Genes are present |
|  | HCN production | + | Genes are present |
|  | Phosphate solubilization | + | Genes are present |
|  | Indole acetic acid (IAA) | + | Genes are present |
|  | Ammonia production | + | Genes are present |
| Hypersensitive response | HR response in the tobacco plant | - | hrp genes are present |

+ sign indicates the presence of the character in the endophytic strain SPT08, whereas -ve represents the absence of the activity

In the genome, *P. aeruginosa* SPT08 has a type III secretion system. To determine its role, hypersensitive response was studied in tobacco leaves by infiltrating 100 μL of saturated bacteria on the lower side of the leaf with a sterile syringe. SPT08 infiltrated area was found to have similar leaf texture and color to *E. coli,* and the sterile water infiltrated area at 48 h. F1C1 exhibited a different leaf discoloration at 12 h, and by 48 h, the leaf became a dry, paper-like texture, suggesting the hypersensitivity reaction (positive control) (Fig. 11). All the observations of the HR test suggest that the endophyte *P. aeruginosa* SPT08 failed to induce the hypersensitive response in the tobacco plant.

**Fig. 11.**
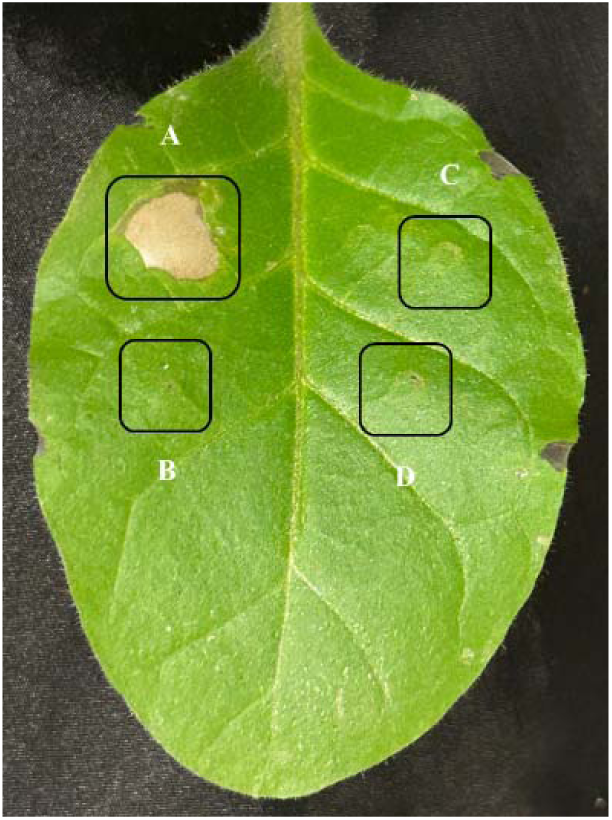
Hypersensitive response assay by *P. aeruginosa* SPT08. Hypersensitive response was assessed by infiltrating the bacterial cell suspension (10^9^ cfu/mL) into the abaxial surface of the tobacco leaves with a sterile syringe. One-month-old tobacco plant leaves were infiltrated with *R. pseudosolanacearum* F1C1 (**A**), *P. aeruginosa* SPT08 (**B**), *E. coli* (**C**), and sterile water (**D**). The picture was taken at 48 h of post-infiltration.

### 3.7 Genomic features, taxonomy, and functional analysis of *P. aeruginosa* SPT08

The whole genome of SPT08 was assembled in 30 contigs with a genome size of 6265489 bp, with a GC content of 66.59% and an N50 value of 440258 bp. The average genome coverage was 430x with 99.66 % completeness. The genome was annotated into 5786 protein-coding sequences through NCBI-PGAP. The genome assembly statistics of SPT08 are listed in Table 5. SPT08 shared an Average Nucleotide Identity (ANI) value of 99.40% with *Pseudomonas aeruginosa* type strain DSM 50071^T^, which confirmed its taxonomic status as per the defined ANI value for the species strains should be equivalent to or more than 95-96% (Kim et al., 2014; Richter and Rosselló-Móra, 2009).

**Table 5.** Genomic characteristics of *P. aeruginosa* SPT08.

| Characteristic | Value |
| --- | --- |
| Genome size (bp) | 6265489 |
| GC content (%) | 66.59 |
| Number of contigs | 30 |
| Average genome coverage (x) | 430x |
| Completeness | 99.66 |
| Contamination | 0.11 |
| N50 value (bp) | 440258 |
| rRNA (5S, 16S, 23S) | 3 |
| tRNA | 52 |
| Protein coding genes (CDSs) | 5786 |

Genomic analysis of SPT08 revealed the presence of genes encoding type I, type II, type III, type V, and type VI protein secretion systems. Further, SPT08 also encodes for type IV pili and flagellum-encoding genes (Supplementary Table S2). SPT08 also encodes for a significant number of biosynthetic gene clusters (BGCs). These included RiPP-like gene clusters, a class of bacteriocins, and multiple Nonribosomal Peptide Synthetase (NRPS) encoding genomic regions, which are attributed to antimicrobial and other important functions. AntiSMASH analysis also predicted beta-lactone and volatile compound haserlactone; these two are well known for antimicrobial activity (Table 6). SPT08 also encodes a hydrogen cyanide gene cluster carrying all three important genes, i.e., *hcnA*, *hcnB*, and *hcnC*. BGC analysis predicted genes encoding three types of siderophores, pyoverdine, pyochelin, and pseudopaline (Table 6, Table 7).

**Table 6:**
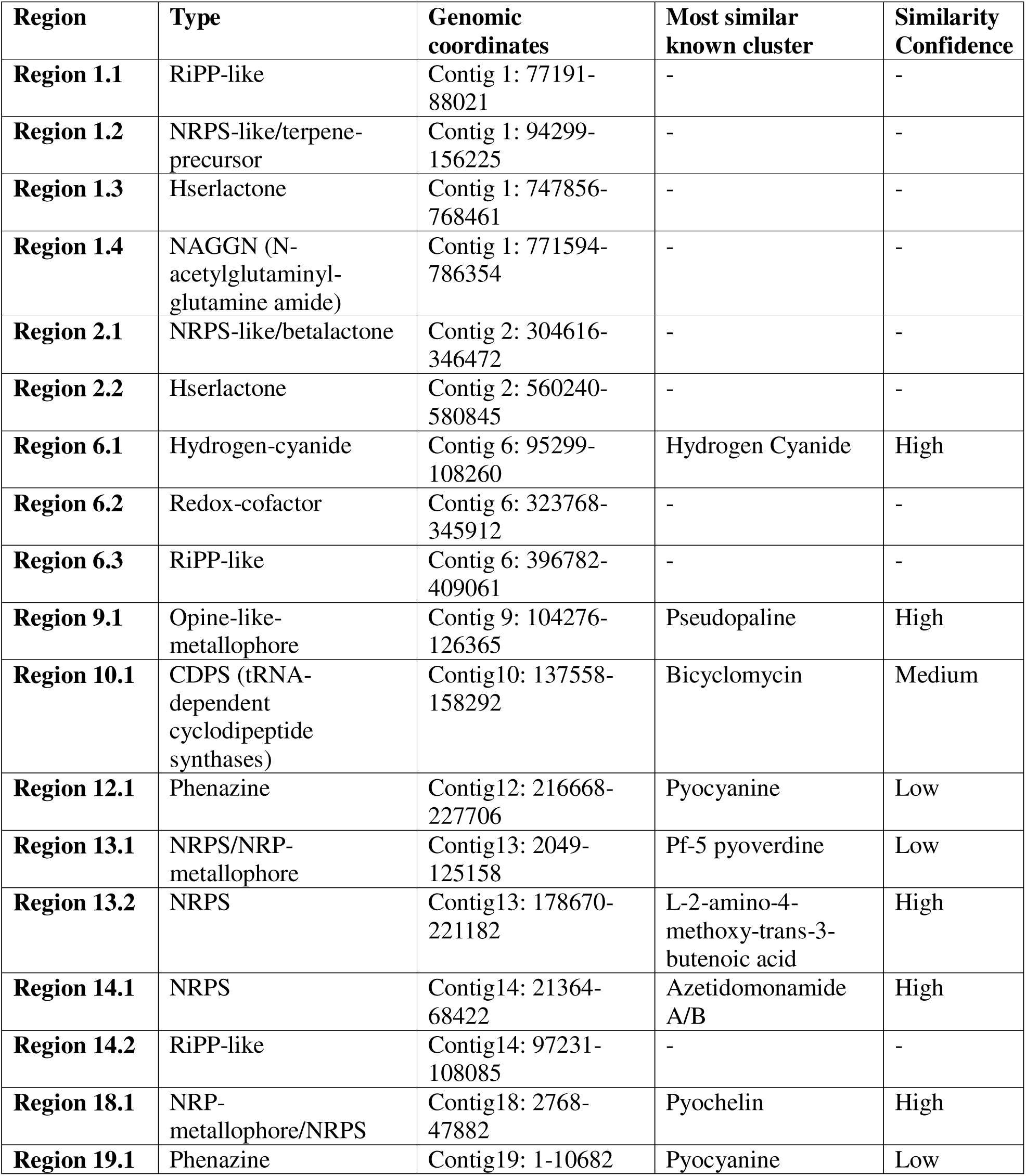
Biosynthetic Gene clusters (BGCs) predicted by antiSMASH in *P. aeruginosa* SPT08, along with their type, genomic coordinates, and most similar known cluster in the databases.

**Table 7.**
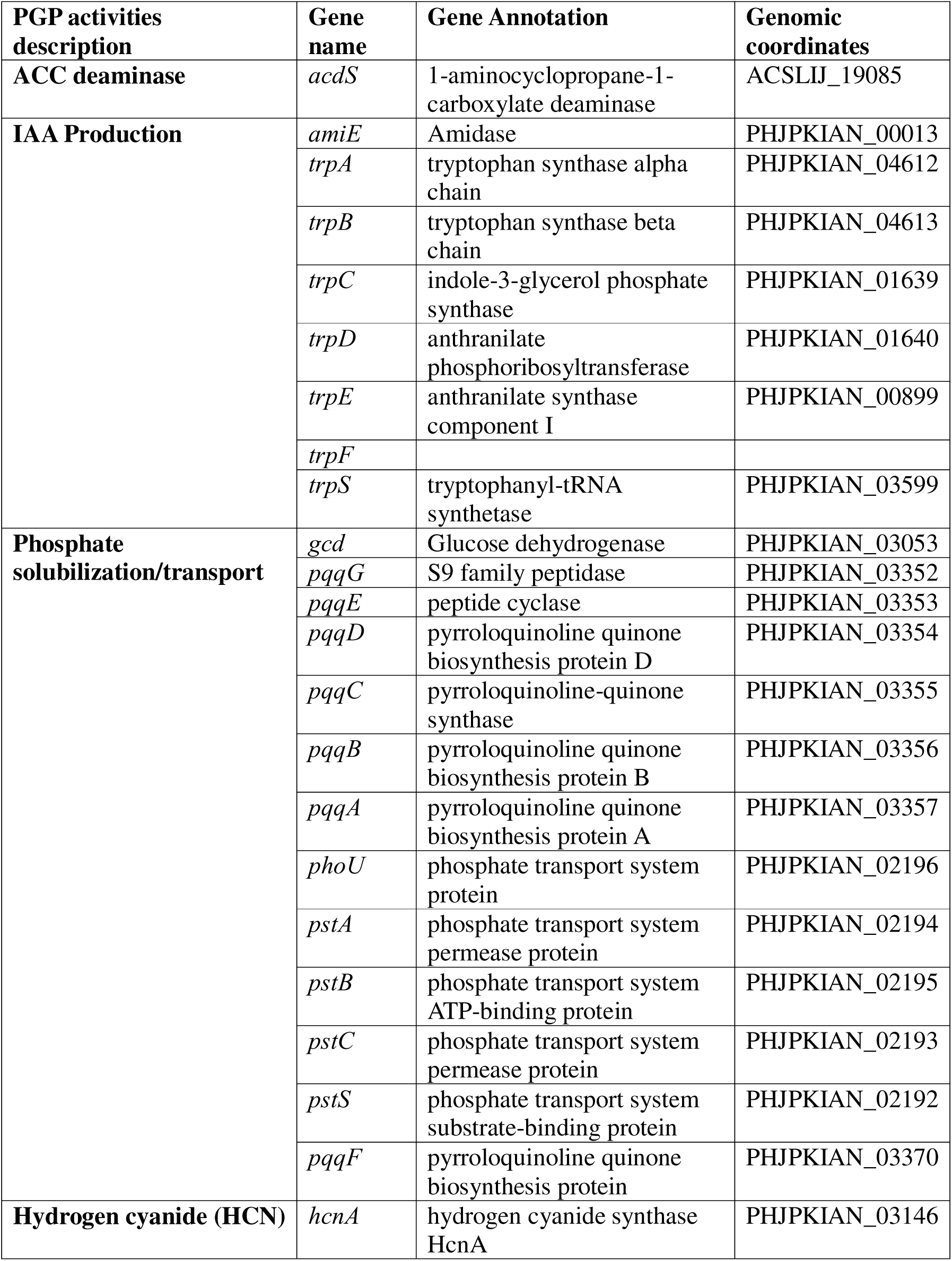
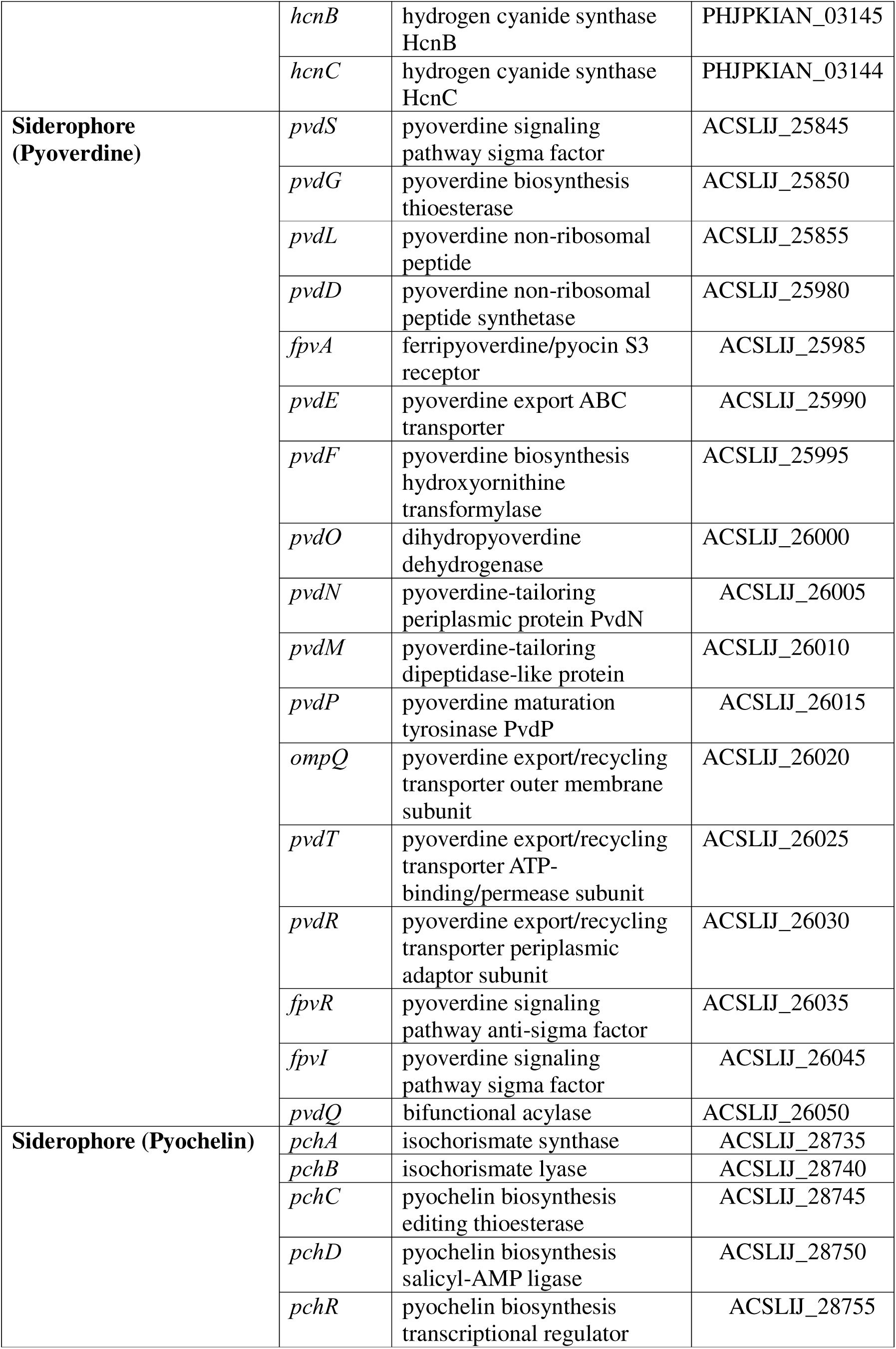
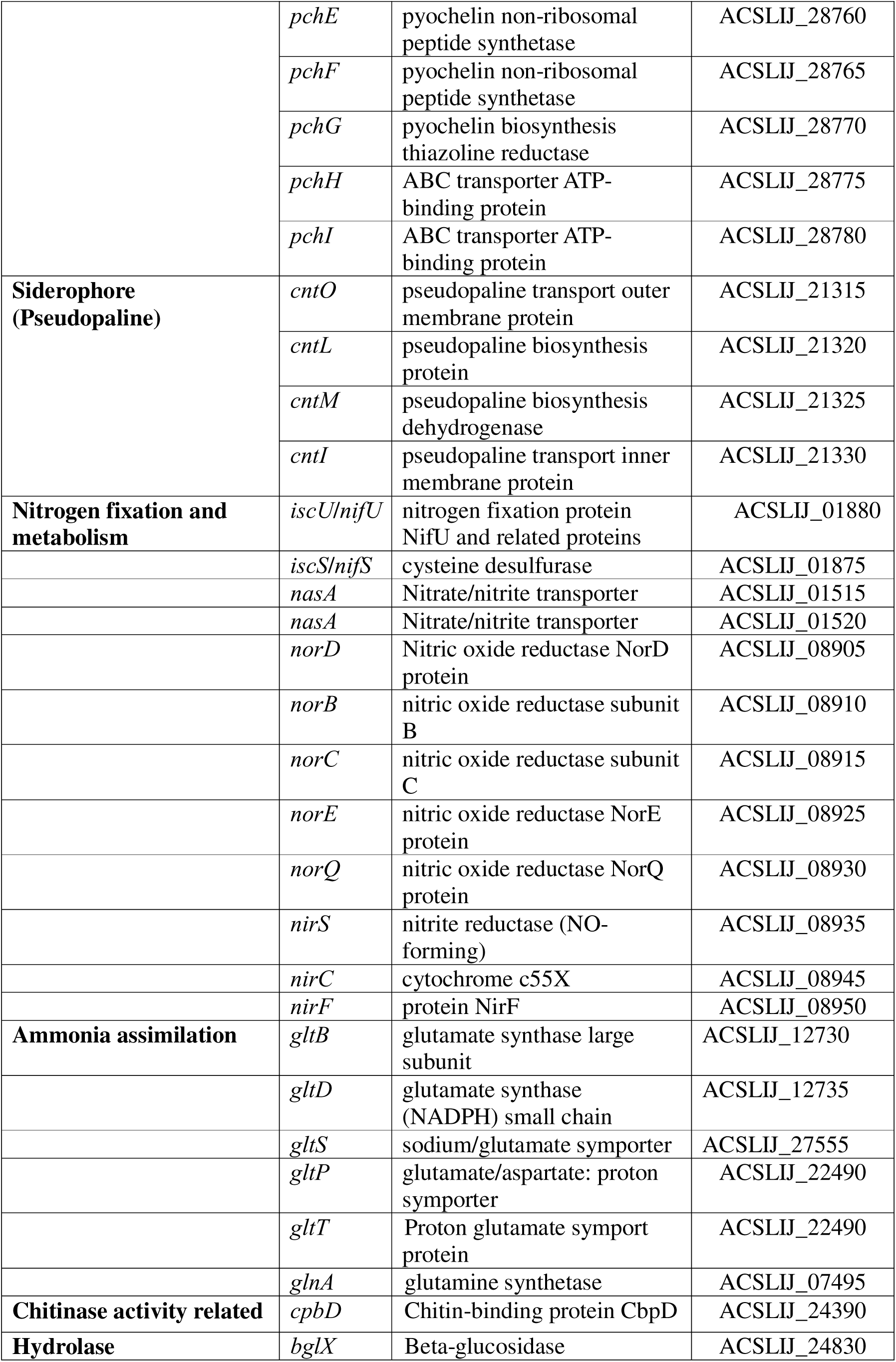
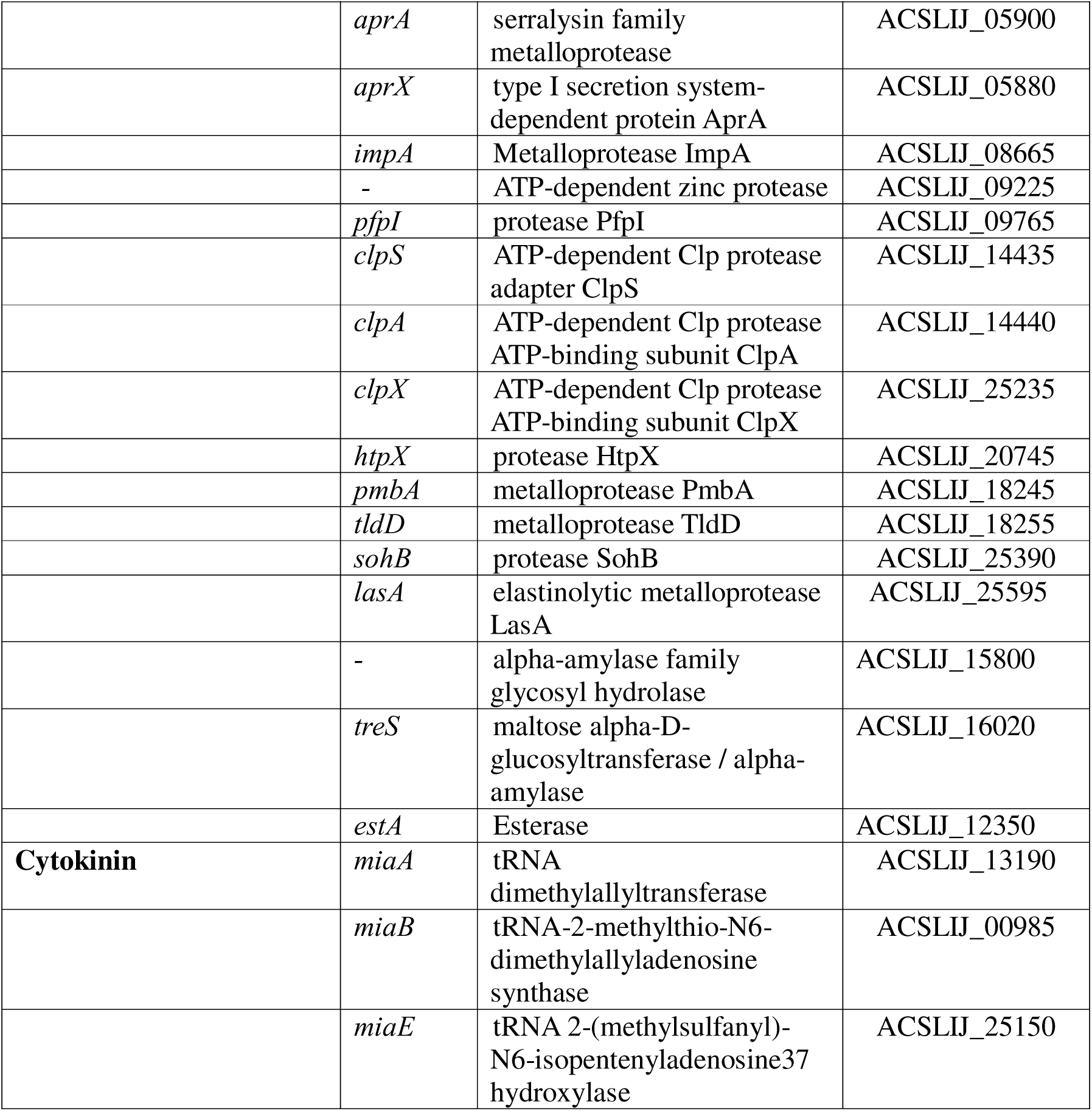
Genes associated with plant growth-promoting (PGP) properties in *P. aeruginosa* SPT08.

*Pseudomonas aeruginosa* SPT08 harbors several genes encoding for PGP properties. These included genes associated with plant growth hormone production, such as IAA, *amiE*, and *trpABCDEFS*, cytokinin, *miaA*, *miaB*, and *miaE,* and ACCD, *acdS*. Further, SPT08 carries a glucose dehydrogenase encoding gene, *gcd,* and a *pqqABCDEG* operon; together, they are involved in phosphate solubilization. It also carries a *pstABCFS* operon involved in phosphate transport. The genome also has a chitin-binding protein encoding gene, *cpbD,* that is known to be associated with chitinase enzymes and could be responsible for anti-fungal properties. The endophyte genome annotation revealed the presence of genes directly or indirectly associated with nitrogen fixation, such as *nifU*, *nifS*, *nasA*, *norBCDQ*, *nirS*, *nirC,* and *nirF*. It also has genes involved in ammonia assimilation. SPT08 carried a large number of cell-wall degrading enzymes encoding beta-glucosidase, metalloproteases, proteases, alpha-amylase, and esterase (Table 7). Interestingly, the genes associated with cellulase activity, such as *celC*, *celE*, and *celA,* were absent in the genome. However, one such gene, *bglX*, was present in the genome of SPT08, encoding for beta-glucosidase.

## 4 Discussion

In this study, several endophytes have been isolated from cotyledon-stage tomato seedlings, of which some of the endophytes exhibit growth inhibition of *Ralstonia pseudosolanacearum*, the causal agent of bacterial wilt in many plants, including tomato. The endophyte *P. aeruginosa* SPT08, which exhibits the maximum growth inhibition activity, has been considered further and demonstrated to protect tomato seedlings as well as grown-up tomato plants against the bacterial wilt. SPT08 also improves the growth of tomato plants and increases their root growth. Using the GFP-tagged SPT08 strain, its colonization has been observed in tomato seedlings as well as in grown-up plants after root inoculation, which proves its true endophyte nature. In addition, SPT08 has several features helpful for plant association, such as the production of extracellular pectinase, exhibiting swimming and twitching motility, and the production of the plant growth hormone, indole acetic acid. The genomic analysis of the SPT08 reveals the presence of homologs of the type III secretion system (T3SS). This secretion system is known to be helpful for the association of bacteria with eukaryotic hosts. The T3SS might be involved in the secretion of various effector proteins, interfering with the cellular functions of the plant host, modulating its physiology and immunity without inducing hypersensitive response and favoring endophytic colonization (Zboralski et al., 2022; Mishra et al., 2021). In *Pseudomonas fluorescens* SBW25, T3SS effectors play an important role in suppressing tomato plant defenses, which facilitates colonization in the host plant without eliciting immune responses (Preston et al., 2001). Remarkably, not all endophytes have a Type III protein secretion system. For example, *Enterobacter roggenkampii* SPT20, an endophyte isolated from the tomato seedlings in our laboratory, does not have a T3SS, which highlights the distinctive feature of SPT08. Genomic analysis confirms the presence of type IV pili in *P. aeruginosa* SPT08, which may contribute to attachment and colonization in the tomato plants. The role of type IV pili has been studied in the endophytic colonization of *Azoarcus* sp. BH72 on rice root surfaces (Böhm et al., 2007).

The systemic colonization of SPT08 in seedlings, as well as in grown-up tomato plants, stimulates several queries about the molecular mechanisms behind the differential adaptation of the endophyte. Many of the unanswered questions include how an endophyte colonizes host plants without causing disease, what makes an endophyte different from a phytopathogen, what determines the host range of an endophyte, and the potential of the endophyte to colonize inside seeds and be inherited from one generation to the next through seeds. In this process of endophyte colonization, some bacterial structural components, such as flagella, pili, and secretory products like lipopolysaccharides, exopolysaccharides, are found to be involved in the establishment of bacteria in the host (Kumar et al., 2020). Proteins that are encoded by genes for adhesion to the host through twitching and swimming motility are frequently used by the endophyte for successful colonization (Mengistu, 2020). The endophyte SPT08 has demonstrated twitching and swimming motility.

The generation time of *P. aeruginosa* SPT08 is ∼ 40 min at 28°C or ∼ 25 min at 37°C, which is much less than that of *R. pseudosolanacearum* F1C1 (generation time ∼2.0 h at 28°C). But we need a 100-fold higher concentration of SPT08 than F1C1 to control the wilt in mixed inoculated tomato plants. We have demonstrated the presence of F1C1 tagged with mCherry in tomato plants that are mix-inoculated with SPT08 tagged with GFP. The colonization of F1C1 in grown-up tomato plants after co-inoculation with SPT08 without causing the wilting of the plants is interesting. This finding aligns with our previous observations of escapee plants, where some of the F1C1 inoculated seedlings remained healthy and did not exhibit wilting symptoms till the experimentation was over, but colonization of F1C1 occurred (Phukan, 2018; Bhuyan et al., 2025). This indicates a greater adaptation ability of F1C1 than SPT08 to survive inside the host plant, that still needs to be investigated in the future.

In grown-up tomato plants, SPT08-mediated increase in plant height and root growth indicates the effect of the endophyte on plant physiology and metabolism. These effects might be contributing towards protection against bacterial wilt in the tomato plant, whose mechanisms may be understood in future research. It is pertinent to note that there is a significant difference between the *in vitro* inhibition of pathogen and the *in-planta* bio-protection. Another endophyte, SPT20 (*Enterobacter roggenkampii*) isolated from tomato seedlings in our laboratory, inhibits F1C1 growth on laboratory medium (Fig. S8), but fails to protect the tomato plant from bacterial wilt in mix-inoculated trials (unpublished result from the laboratory). This evidence indicates that all endophyte inhibiting a pathogen *in vitro* will not necessarily be capable of protecting against the disease. Moreover, experimental evidence with eggplants being mix-inoculated with SPT08 and F1C1, providing only partial protection (as some eggplant plants developed symptoms of bacterial wilt), indicates host-specific limitation of SPT08 protection ability. These varying consequences across the two different host plants indicate that the protection mechanism of SPT08 involves a complex interplay between the endophyte, F1C1, and the host plant.

*P. aeruginosa* SPT08 causes a significant alteration in the root architecture of tomato plants through various plant growth-promoting traits, which ultimately enhances host vitality and controls *R. pseudosolanacearum* F1C1 infection. SPT08 produces indole-3-acetic acid, an important phytohormone for the growth of root and shoot tissues. Auxin, a key plant hormone, plays a significant role in root modification and regulates root growth through nutrient uptake and the activation of immune or defense responses. According to Glick et al. (2003), *Pseudomonas* spp. enhances root branching by secreting auxin and reducing ethylene via ACC deaminase. SPT08 may modulate the plant auxin biosynthesis pathway or interact with other signaling molecules to fine-tune root architecture. Genome analysis confirms the presence of auxin biosynthesis genes in SPT08 that may be involved in the growth of root architecture in tomato plants. Apart from auxin, microbial metabolites also contribute to plant growth by altering the root architecture (Lugtenberg and Kamilova, 2009). Auxin-like small cyclic peptides, such as diketopiperazine, and volatile molecules are produced by many endophytic bacteria, which are shown to be involved in the structural modification of plant roots. Likewise, metabolites like acetoin and 2,3-butanediol secreted by *Pseudomonas* PS01 are involved in plant growth regulation and alter root structure. *P. aeruginosa* SPT08 genome contains genes which are involved in the synthesis of acetoin, 2,3-butanediol, and bicyclomycin (2,5-diketopiperazines). Plant root growth promotion by SPT08 and change in root structure might be independent of bacterial auxin or by a combination of different volatile and non-volatile compounds. Many bacteria, such as *Pseudomonas* sp., *Bacillus* sp., *P. fluorescens*, and *B. subtilis*, as well as *A. brasilense*, are reported to be involved in promoting plant growth and influencing root morphology (Glick et al., 2003; Bashan and de-Bashan, 2010). The mechanism of endophyte-induced root growth is still unknown; however, the auxin or auxin-like molecule derived from microbes affects the root physiology and function (Overvoorde et al., 2010; Bouzroud et al., 2018; Sukumar et al., 2013; Ortíz-Castro et al., 2009). Moreover, the mechanism of plant growth includes the mobilization of nutrients, such as phosphate, iron, etc., which are involved in root development. The endophyte SPT08 has both phosphate-solubilizing and siderophore-producing activities. Phosphorus is mainly required for the processes involved in strong root system development and increases nutrient and water absorption by the host plant (Khan et al., 2023). Among the most potent phosphate solubilizers are strains belonging to the bacterial genera such as *Pseudomonas*, *Bacillus*, and *Rhizobium*, which are reported as potent phosphate solubilizers having a role in root growth (Rodrı guez and Fraga, 1999). Siderophores are involved in increasing the pathogen resistance by solubilizing the insoluble iron and suppressing the pathogen growth by limiting the iron source, enabling the endophyte to establish the ecological niche in and around the rhizosphere effectively (Egamberdieva et al., 2023). By solubilizing insoluble iron and limiting its availability to *R. pseudosolanacearum*, SPT08 enhances plant health and establishes a competitive ecological niche in the rhizosphere. Furthermore, SPT08 produces hydrogen cyanide and ammonia. HCN indirectly controls the bacterial growth of phytopathogens, whereas ammonia facilitates the absorption of ammonium by the host plant (Kommedahl, 1984; Dordas, 2008). All these processes are involved in increasing the availability of nutrients, water uptake, and changing the root architecture of the plant.

SPT08 produces several hydrolytic enzymes, such as pectinase, protease, and amylase, except cellulase, which is confirmed after genomic analysis of the bacteria. This enzymatic characterization helps the endophyte to strategize its colonization. The absence of cellulase helps to maintain the structural integrity of the host cells, and the presence of pectinase helps in the target degradation of specific components of the plant cell wall, enabling the bacteria to enter the host cells. Pectinase activity in *Pseudomonas aeruginosa* has not been reported widely except in a few environmental isolates (Sohail and Latif, 2016), which might be indicative of plant adaptation. SPT08 appears to employ these hydrolytic enzymes at favorable sites to invade host plant cells. The protease enzyme of SPT08 may be involved in the degradation of protein or proteolytic enzymes secreted by the pathogen *R. pseudosolanacearum*. Endophytes produce extracellular amylase that might be involved in the invasion of host plants (Pathak et al., 2022; Reinhold-Hurek and Hurek, 1998; Elbeltagy et al., 2000). Amylase activity of SPT08, demonstrated by starch hydrolysis on agar plates, likely aids in breaking down complex carbohydrates in the rhizosphere or plant tissues. This provides energy for colonization and may facilitate root invasion by modifying local nutrient availability, which may also be involved in root growth (Agustiani et al., 2023; Huang et al., 2022). However, an elaborate exploration is required of the enzymatic secretion of endophytic bacteria and their role in the invasion of the host plant and colonization efficiency.

Despite showcasing remarkable biocontrol activity against bacterial wilt and significant plant growth-promoting abilities, the possibility of *P. aeruginosa* being an opportunistic pathogen cannot be eliminated without future analysis of comparative genomics of SPT08 with nosocomial *P. aeruginosa* strains. Studies including characterization of promising latent virulence factors, assessing possible agricultural risks, will help us better understand if the benefits provided by SPT08 are due to genomic adaptation that controls wilting or is just a phenotypic response to environmental factors.

## 5 Conclusion

Endophytes are one type of plant probiotics that keep the plant healthy. Endophyte *Pseudomonas aeruginosa* SPT08 has a significant role in crop protection properties against wilt disease and promotes plant growth. SPT08 also enhances the plant root growth and plant height; the mechanism of inhibition, as well as root growth, is under investigation. The strain may be used as a biocontrol agent to protect plants and minimize the use of chemical fertilizers for sustainable agriculture. This is the first report of an endophyte from tomato seedlings that protects the host plant against bacterial wilt caused by *R. pseudosolanacearum*.

## 6 Future perspective

We will be looking forward to the antifungal and insecticidal activity of the endophyte *P. aeruginosa* SPT08 against various plant pathogens and the crop protection ability of the endophyte in other solanaceous crops. However, further studies will be required for field application by directly applying the bacterial cell suspension or in the form of a formulation to explain the usability of the bacteria as a microbial biocontrol agent. How does the endophyte provide benefits to the whole plant, and what are the molecular mechanisms of root growth? The root morphology change might induce xylogenesis in tomato plants, leading to bacterial wilt resistance? These are some questions that will be addressed in future research.

## Supplementary data

Table S1. Plant growth parameters of grown-up plants of two independent experiments performed under the greenhouse

Table S2. Type IV pili and flagellum-encoding genes in *P. aeruginosa* SPT08

Fig. S1. Morphological characterization of *Pseudomonas aeruginosa* SPT08

Fig. S2. Motility of endophyte *P. aeruginosa* SPT08

Fig. S3. *Pseudomonas aeruginosa* SPT08 biocontrol activity against bacterial wilt in grown-up tomato plants under greenhouse conditions

Fig. S4. *Pseudomonas aeruginosa* SPT08 promotes the growth of the tomato plant height

Fig. S5. *Pseudomonas aeruginosa* SPT08 impact on the tomato root growth

Fig. S6. Increase in root fresh weight and dry weight by endophyte *P. aeruginosa* SPT08

Fig. S7. Longitudinal root length of tomato plants inoculated with *P. aeruginosa* SPT08

Fig. S8. *Ralstonia pseudosolanacearum* F1C1 growth inhibition by *Enterobacter roggenkampii* SPT20

## Acknowledgements

We thank Prof. Kirankumar S. Mysore from Oklahoma State University, UAS, and the former Professor of Plant Biology Division at The Samuel Roberts Noble Foundation, USA, for his kind gift of the pDSK-GFPuv plasmid vector. We are grateful to Prof. Marc Valls for his kind gift of the mCherry plasmid vector. The authors thank Dr. Rahul Kumar and Dr. Ruksana Aziz from SKR Lab, who initially started the antibacterial activity assay in the laboratory. SJG and SB are thankful for the JRF fellowship from the DBT, GoI, New Delhi, under grant (BT/PR41637/NER/95/1753/2021) awarded to Suvendra Kumar Ray. RR is thankful for the JRF/SRF fellowship from the CSIR, GoI, New Delhi. PLS appreciates the JRF/SRF fellowship from the DBT, GoI, New Delhi, granted under (BT/403/NE/U-Excel/2013) awarded to Suvendra Kumar Ray. LD expresses gratitude to DST, GoI, New Delhi, for the INSPIRE-SRF fellowship. SB is thankful for the JRF/SRF fellowship from the UGC-NFSC, GoI, New Delhi. SK and MJ acknowledge the JRF fellowship from the UGC-CSIR, GoI, New Delhi.

## Author contributions

**SJG** performed the experiments, analyzed the data, prepared the draft manuscript, reviewed, and edited. **RR** did the genomic analysis, reviewed, and edited. **SB** and **LD** helped in the greenhouse experiments, performed formal data analysis, reviewed, and edited the article**. SK, SB, and MJ** performed formal data analysis, reviewed the manuscript. **PLS** and **NA** performed the biocontrol activity using *Pseudomonas* sp. in seedlings, formal data analysis, and reviewed the manuscript. **SKR** and **PBP** supervised, reviewed, and edited, and conceptualization. All the authors have read the manuscript and declare no conflict of interest.

## Data availability

Genome data of *Pseudomonas aeruginosa* SPT08 has been submitted to the NCBI with Accession No. JBPQBA000000000. All the other data generated during this study are included in the manuscript and supplementary files.

---

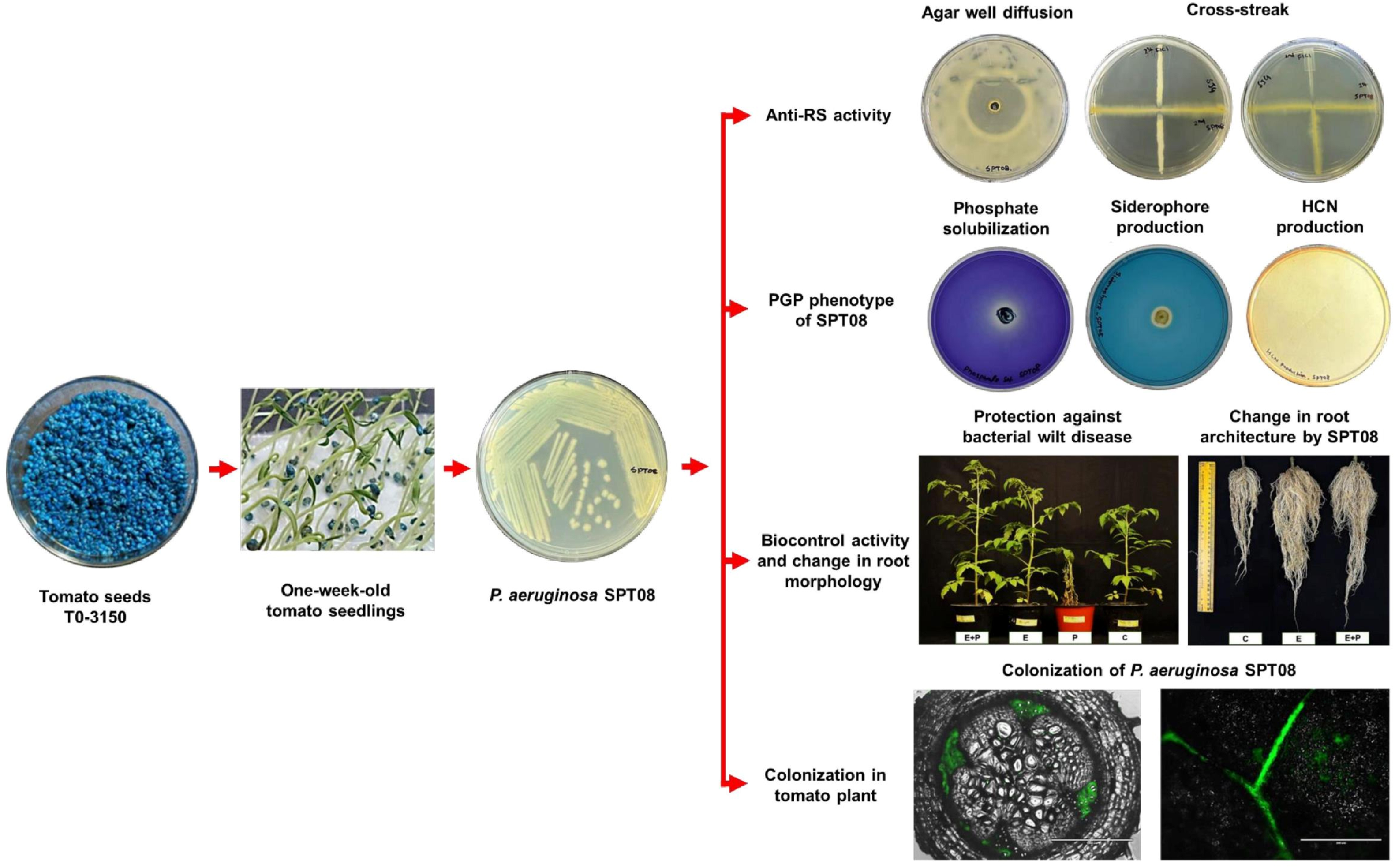

